# A dual function of TMEM70 in OXPHOS: assembly of complexes I and V

**DOI:** 10.1101/697185

**Authors:** Laura Sánchez-Caballero, Dei M. Elurbe, Fabian Baertling, Sergio Guerrero-Castillo, Mariel van den Brand, Joeri van Strien, Teunis J. P. van Dam, Richard Rodenburg, Ulrich Brandt, Martijn A. Huynen, Leo G.J. Nijtmans

## Abstract

Protein complexes from the oxidative phosphorylation (OXPHOS) system are assembled with the help of proteins called assembly factors. We here delineate the function of the inner mitochondrial membrane protein TMEM70, in which mutations have been linked to OXPHOS deficiencies, using a combination of BioID, complexome profiling and coevolution analyses. TMEM70 interacts with complex I and V and for both complexes the loss of TMEM70 results in the accumulation of an assembly intermediate followed by a reduction of the next assembly intermediate in the pathway. This indicates that TMEM70 has a role in the stability of membrane-bound subassemblies or in the membrane recruitment of subunits into the forming complex. Independent evidence for a role of TMEM70 in OXPHOS assembly comes from evolutionary analyses. The TMEM70/TMEM186/TMEM223 protein family, of which we show that TMEM186 and TMEM223 are mitochondrial in human as well, only occurs in species with OXPHOS complexes. Our results validate the use of combining complexomics with BioID and evolutionary analyses in elucidating congenital defects in protein complex assembly.

## Introduction

The oxidative phosphorylation (OXPHOS) system, situated in the mitochondrial inner membrane, is composed of five enzyme complexes (I–V) and two electron carriers, coenzyme Q and cytochrome *c*^1^. The first four complexes form the electron transport chain (ETC) that couples the transfer of electrons from NADH and Succinate to oxygen, to the transfer of protons across the membrane. The thus generated proton gradient is used by complex V (CV) or ATP synthase to generate adenosine triphosphate (ATP)^2^. From the five enzymes of the OXPHOS system, the first and the last, i.e., complex I (CI) or NADH:ubiquinone oxidoreductase and CV are particularly relevant to this study. Mammalian CI is a 45 subunit L-shaped complex, seven subunits of which are mitochondrially encoded, (MT-ND1, MT-ND2, MT-ND3, MT-ND4, MT-ND4L, MT-ND5 and MT-ND6) with a lipophilic arm integrated into the inner mitochondrial membrane and a hydrophilic peripheral arm jutting out into the mitochondrial matrix^3^. The enzyme can be subdivided into three different functional modules^4^ that are assembled separately^5,6^: the P-module (proton translocation) that represents the lipophilic membrane arm, the N-module (NADH dehydrogenase) and the Q-module (coenzyme Q reduction). Mammalian CV is an enzyme complex composed of 18 subunits (including the regulatory protein IF_1_), two of which are mitochondrially encoded (MT-ATP6 and MT-ATP8)^7^. It comprises two modules: the globular F_1_ domain (subunits α, β, γ, δ and ε) that contains the catalytic region, and the membrane-embedded F_O_ domain, that contains the rotary motor. The F_1_ module is connected to F_O_ (subunits DAPIT, 6.8PL, a, A6L, e, f, g and c) by two stalks, one central, with subunits γ, δ, ε and one peripheral that consists of the oligomycin sensitivity conferral protein (OSCP) and subunits b, d, and F_6_. The central stalk and the c-subunits octamer establish the rotor of the fully assembled enzyme^8^.

For the optimal assembly of the OXPHOS complexes extra proteins are needed: the so-called assembly factors, which assist in the assembly of a complex but are not part of the final complex. Assembly factors have been discovered by genetic and experimental approaches, often in combination with bioinformatic analyses^9^. Those analyses have exploited that the assembly factors are often part of large protein families^10^. They show conservation of molecular function and co-evolve with CI itself^11^. This is e.g. the case for the CI assembly factor IND1, that was predicted to play a role in the assembly of Fe-S clusters in CI via the role of its cytoplasmic homologs in the assembly of other Fe-S proteins and via its co-evolution with CI proteins^12^. However, the exact molecular function of many assembly factors has not been elucidated.

From all the complexes that are part of the OXPHOS system, the one with the lowest number of known assembly factors is CV. Its known assembly factors are ATPAF1 and ATPAF2^13^, which both interact with the F_1_ module, and TMEM70^14^. TMEM70 is a transmembrane protein that localizes in the inner membrane of the mitochondria^15,16^. It contains two transmembrane regions that form a hairpin structure of which the N- and C-termini are located in the mitochondrial matrix^17^. Mutations in *TMEM70* have been reported to severely diminish the content of CV in a large cohort of patients^14,18-26^, and of all nuclear encoded proteins affecting CV, TMEM70 is the most commonly mutated in disease^27^. Moreover, *Tmem70* knockout mouse embryos show stalled F_1_ module^28^. These observations led to the hypothesis that TMEM70 is a CV assembly factor. However, there is no evidence of its direct interaction with CV proteins, and its specific role in CV assembly remains unclear^16,26^.

Interestingly, defects in TMEM70 have not exclusively reported deleterious effects on CV but also that combined with less severe CI deficiency^22^, a combined OXPHOS deficiency^29^ or even an isolated CI deficiency^20^. Furthermore, TMEM70 has been shown to co-migrate with an assembly intermediate that forms part of CI^6^, suggesting it might form part of its assembly process.

The present study aims to elucidate the full role of TMEM70 in the OXPHOS system using standard biochemical techniques combined with two novel techniques: complexome profiling and the BioID proximity-dependent labelling assay, together with an in-silico approach to detect TMEM70 homologs and reconstruct the co-evolution of TMEM70 with other mitochondrial proteins.

## Results

### BirA* tagged TMEM70 biotinylates complex I, complex V and the small subunit of the ribosome

To study whether there is a direct interaction of TMEM70 with components of CI and CV and thus link its absence to both OXPHOS deficiencies, or whether it interacts with any other mitochondrial protein complex, we used HEK293 Flp-In T-Rex293 cell lines engineered to express BirA*- tagged TMEM70 in a doxycycline-inducible manner (BioID). In this experiment, designed to biotinylate proteins that occur in close proximity to our protein of interest, we detected 538 proteins (Supplementary Table S1). From those, we obtained a list of 135 putative interaction partners of TMEM70 based on proteins showing a significant increase in biotin positive – doxycycline positive conditions compared to biotin positive – doxycycline negative conditions and biotin negative – doxycycline positive conditions (Supplementary Fig. S1, Supplementary Table S2). The mitochondrial proteins biotinylated by BirA*-TMEM70 (n = 102) were significantly enriched for proteins involved in CI – including assembly factors NDUFAF1, NDUFAF2, NDUFAF4 and NDUFAF5 -– and CV, as well as the small subunit of the mitochondrial ribosome (analysis done with DAVID^30^, Table 1, Supplementary Table S3).

**Table 1.** Enriched clusters obtained from potential mitochondrial interactors of TMEM70.

| Category | Term | GO | Cluster | FDR |
| --- | --- | --- | --- | --- |
| GOTERM_BP_DIRECT | mitochondrial respiratory chain complex I assembly | GO:0032981 | 1 | 6.53e-11 |
| GOTERM_BP_DIRECT | mitochondrial electron transport, NADH to ubiquinone | GO:0006120 | 1 | 4.73e-6 |
| GOTERM_CC_DIRECT | mitochondrial respiratory chain complex I | GO:0005747 | 1 | 8.03e-5 |
| GOTERM_MF_DIRECT | NADH dehydrogenase (ubiquinone) activity | GO:0008137 | 1 | 9.56e-5 |
| GOTERM_BP_DIRECT | mitochondrial translational elongation | GO:0070125 | 2 | 4.02e-4 |
| GOTERM_CC_DIRECT | Mitochondrial small ribosomal subunit | GO:0005763 | 2 | 1.15e-3 |
| GOTERM_BP_DIRECT | Mitochondrial translation termination | GO:0070126 | 2 | 6.17e-3 |
| GOTERM_CC_DIRECT | Mitochondrial proton-transporting ATP synthase complex | GO:0005753 | 3 | 1.85e-2 |
| GOTERM_CC_DIRECT | Mitochondrial ATP synthesis coupled proton transport | GO:0042776 | 3 | 2.60e-2 |

### A comprehensive assessment of the *TMEM70* knockout effect on the OXPHOS system

An interaction of TMEM70 with CI and CV, as we observed in the BioID results, might explain why patients harbouring *TMEM70* mutations have CI and CV deficiencies^20,22,29^. To obtain a detailed view on how the absence of TMEM70 affects those mitochondrial protein complexes, we performed complexome profiling on a *TMEM70* knockout cell line and on controls. We compared two HAP1 wild-type cell lines with one mutant cell line that had a 32bp deletion in exon 1 of the *TMEM70* gene, and performed a biological replicate by again enriching mitochondria from these cell lines and repeating the complexome profiling. The deletion in exon 1 of *TMEM70* causes a frameshift and a premature stop codon in the cDNA, with no detectable presence of the protein by MS/MS (Table 2). Complexome profiling, which consists of a BN-PAGE followed by mass spectrometry, allows us to see the majority of the proteins belonging to each of the native complexes and their distribution over the various assembly intermediates^31,32^. In the six complexome profiles, four controls and two *TMEM70* knockouts, we detected in total 3766 proteins, of which 814 are annotated as mitochondrial in MitoCarta 2.0^33^. The profiles were normalized such that the sum of the intensities of the mitochondrial proteins between the samples were equal, and the profiles were aligned with COPAL^34^ to allow comparison between the replicates (see Supplementary Table S4 for the complete results).

**Table 2.**
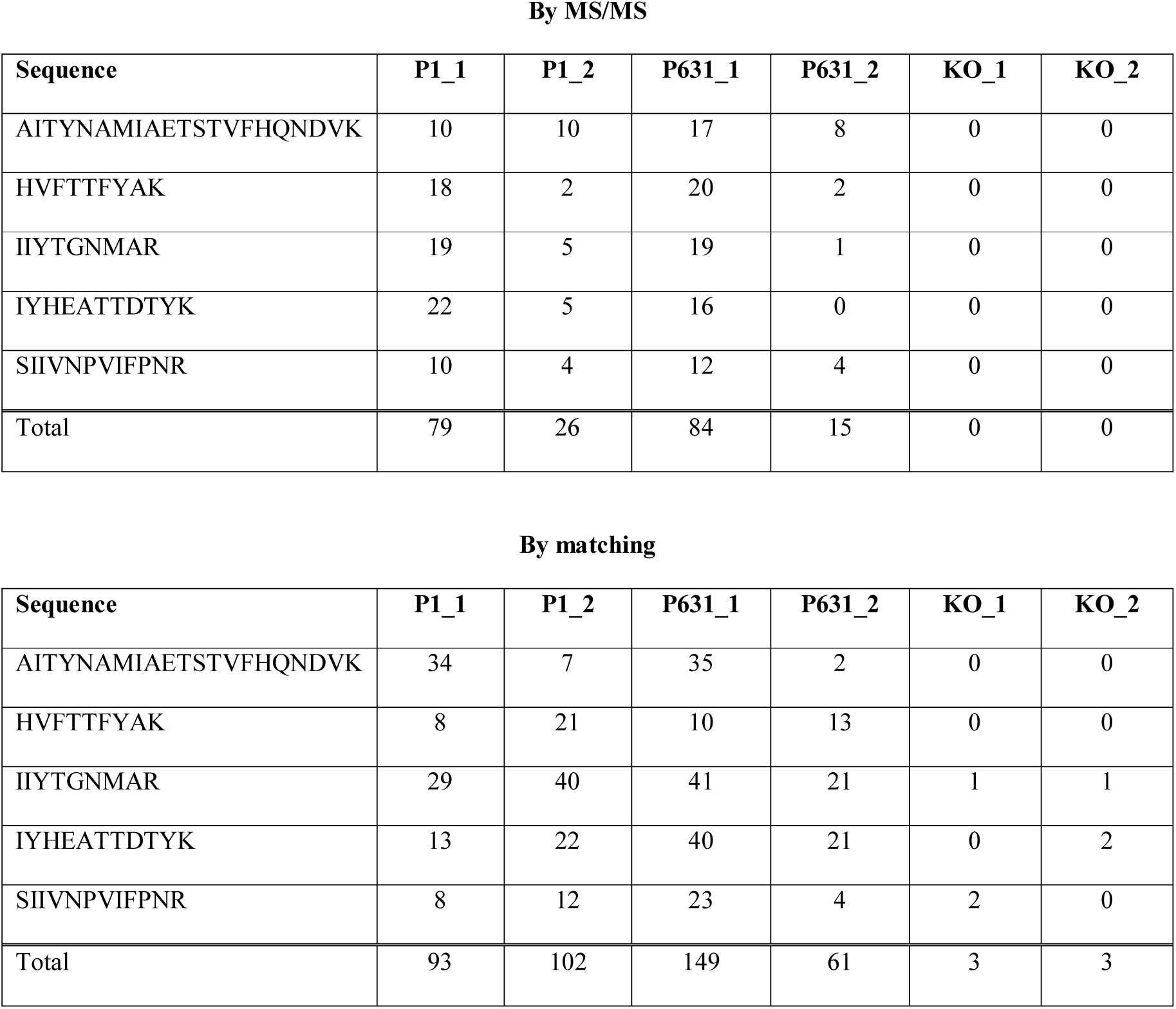
Peptides assigned to TMEM70 obtained in all 6 samples by MS/MS and by matching.

By MS/MS
| Sequence | P1_1 | P1_2 | P631_1 | P631_2 | KO_1 | KO_2 |
| --- | --- | --- | --- | --- | --- | --- |
| AITYNAMIAETSTVFHQNDVK | 10 | 10 | 17 | 8 | 0 | 0 |
| HVFTTFYAK | 18 | 2 | 20 | 2 | 0 | 0 |
| IITYGNMAR | 19 | 5 | 19 | 1 | 0 | 0 |
| IYHEATTDITYK | 22 | 5 | 16 | 0 | 0 | 0 |
| SIIVNPVIFPNR | 10 | 4 | 12 | 4 | 0 | 0 |
| Total | 79 | 26 | 84 | 15 | 0 | 0 |

| Sequence | P1_1 | P1_2 | P631_1 | P631_2 | KO_1 | KO_2 |
| --- | --- | --- | --- | --- | --- | --- |
| AITYNAMIAETSTVFHQNDVK | 34 | 7 | 35 | 2 | 0 | 0 |
| HVFTTFYAK | 8 | 21 | 10 | 13 | 0 | 0 |
| IITYGNMAR | 29 | 40 | 41 | 21 | 1 | 1 |
| IYHEATTDITYK | 13 | 22 | 40 | 21 | 0 | 2 |
| SIIVNPVIFPNR | 8 | 12 | 23 | 4 | 2 | 0 |
| Total | 93 | 102 | 149 | 61 | 3 | 3 |

In order to assess the overall effects caused by the lack of TMEM70 on OXPHOS, and to confirm its interactions with complexes I and V suggested by our BioID results (Table 1) as well as by previous studies^6,22^, we measured the presence of the subunits belonging to the fully formed OXPHOS complexes (see Methods). We observed a significant decrease in the abundance of the subunits belonging to CI and CV and small, non-significant increases in the other three OXPHOS complexes (Fig. 1a; *p* values: CI – 1.4e-12 (median ∼60% compared to controls), CII – 0.125, CIII – 0.322, CIV – 0.204, CV – 1.5e-5 (median ∼30% compared to controls), Wilcoxon signed-rank test, Supplementary Fig. S2). We also tested their enzymatic activities by spectrophotometric analysis and, accordingly, results showed a 30% decrease in CI activity (*p* = 0.156) and a 70% decrease and CV activity (*p* = 0.009), Fig. 1b), whereas the other OXPHOS complexes showed no apparent change in activity. Furthermore, we also observed a reduction in fully formed complexes I and V in BN-PAGE/immunoblotting analysis in KO conditions compared to controls (Supplementary Fig. S3).

**Figure 1.**
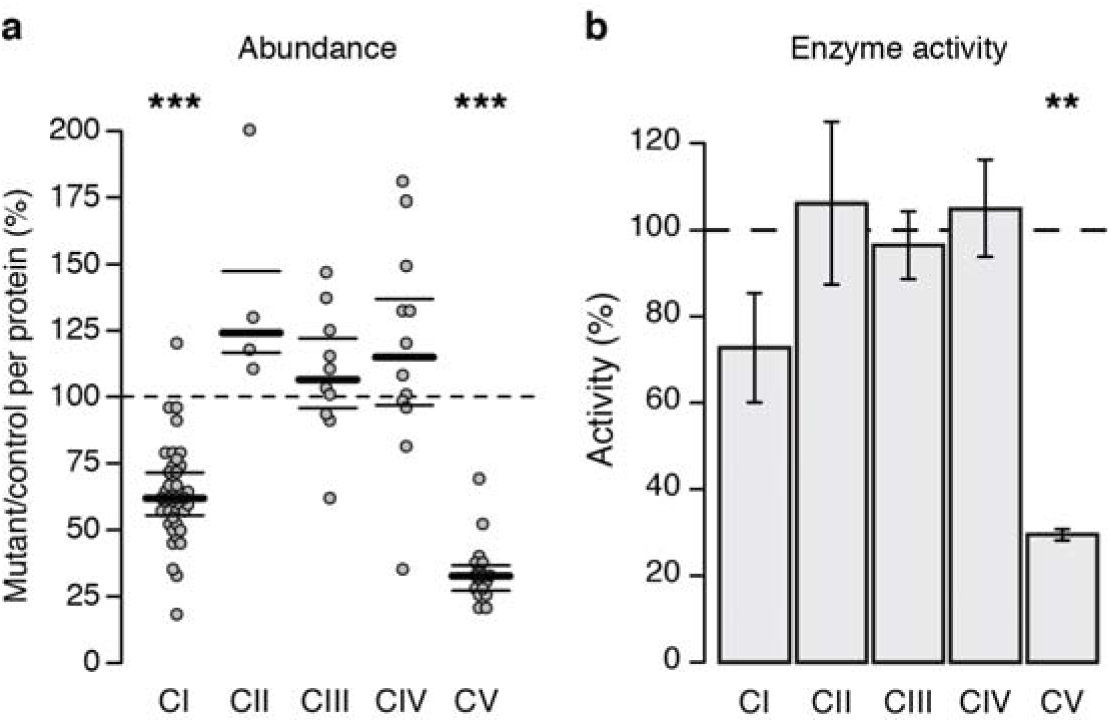
Lack of TMEM70 has an effect in CI and CV. **a** Ratios between mutants and controls measured per subunit and grouped by complex. CI and CV show an apparent reduction of ∼40% and 70% respectively. **b** Enzymatic activities shown by TMEM70 KO with respect to the parental control cell line. Complexes I and V show a decrease, although statistically significant only for the latter (*p* = 0.156, 0.8, 0.51, 0.577 and 0.009, for complexes I, II, III, IV and V, respectively; n = 3). Error bars represent the standard deviation.

### Impaired assembly of complex I

We first focused on the differences in the assembly process of CI between the *TMEM70* wild-type and in *TMEM70* knockout HAP1 cells. As mentioned above, there was a reduction of fully formed CI in cells lacking TMEM70 (Fig. 1a), suggesting a possible CI assembly defect. CI assembly proceeds via pre-assembled intermediates that later on interact, until the fully formed and active enzyme is made^5^. When analysing the assembly steps of CI under *TMEM70* KO conditions in the complexomics results, we observed a clear and significant accumulation of the Q/P_P_-a intermediate (Fig. 2 and Supplementary Fig. S4; median ratio = 4.14; *p* = 4.89e-4) relative to controls, consistent with the accumulation observed in two-dimensional BN-PAGE/SDS-PAGE for NDUFS3 intermediates (Supplementary Fig. S5). At this stage, the Q-module formed by NDUFS2, NDUFS3, NDUFS7, NDUFS8 and NDUFA5 and the assembly factors NDUFAF3 and NDUFAF4 is anchored to the membrane with proteins belonging to the proximal P-module, TIMMDC1 and MT-ND1^6^. Following the formation of this intermediate, and based on one of the previously described assembly pathways of complex I (Fig. 2), the remaining subunits of the proximal module (P_P_-b), which do not show any significant changes while forming part of this intermediate (Fig. 2 and Supplementary Fig. S4; median ratio = 0.94; *p* = 0.102), are added to form the intermediate Q/P_P_ that, in the TMEM70 depleted cells, also accumulates relative to the controls (Fig. 2 and Supplementary Fig. S4; median ratio = 2.41; *p* = 2.6e-3). The most distal part of the enzyme, the P_D_ intermediate, is composed of two different intermediates: P_D_-a and P_D_-b. TMEM70 has been observed to co-migrate with this distal membrane part (P_D_) of complex I, and within this distal part, with the intermediate P_D_-a, that is just next to the proximal membrane part of the complex^6^. With respect to the P_D_ part of the enzyme, the intermediate P_D_-b showed a minor but significant accumulation (Fig. 2 and Supplementary Fig. S4; median ratio = 2.18; *p* = 0.016), while no accumulation was observed for P_D_-a (median ratio = 1.04; *p* = 0.383). These results are as well compatible with the alternative assembly pathway of complex I (Supplementary Fig. S6), which also shows a significant decrease of the intermediates that follow after the incorporation of P_D_-a (P_P_-b/P_D_-a, Supplementary Fig. S6 and Supplementary Fig. S4; median ratio = 0.64; *p* = 0.004). Finally, both assembly pathways converge with the formation of intermediate Q/P, which is also reduced in the absence of TMEM70 (Fig. 2, Supplementary Fig. S6 and Supplementary Fig. S4; median ratio = 0.70; *p* = 1.9e-7).

**Figure 2.**
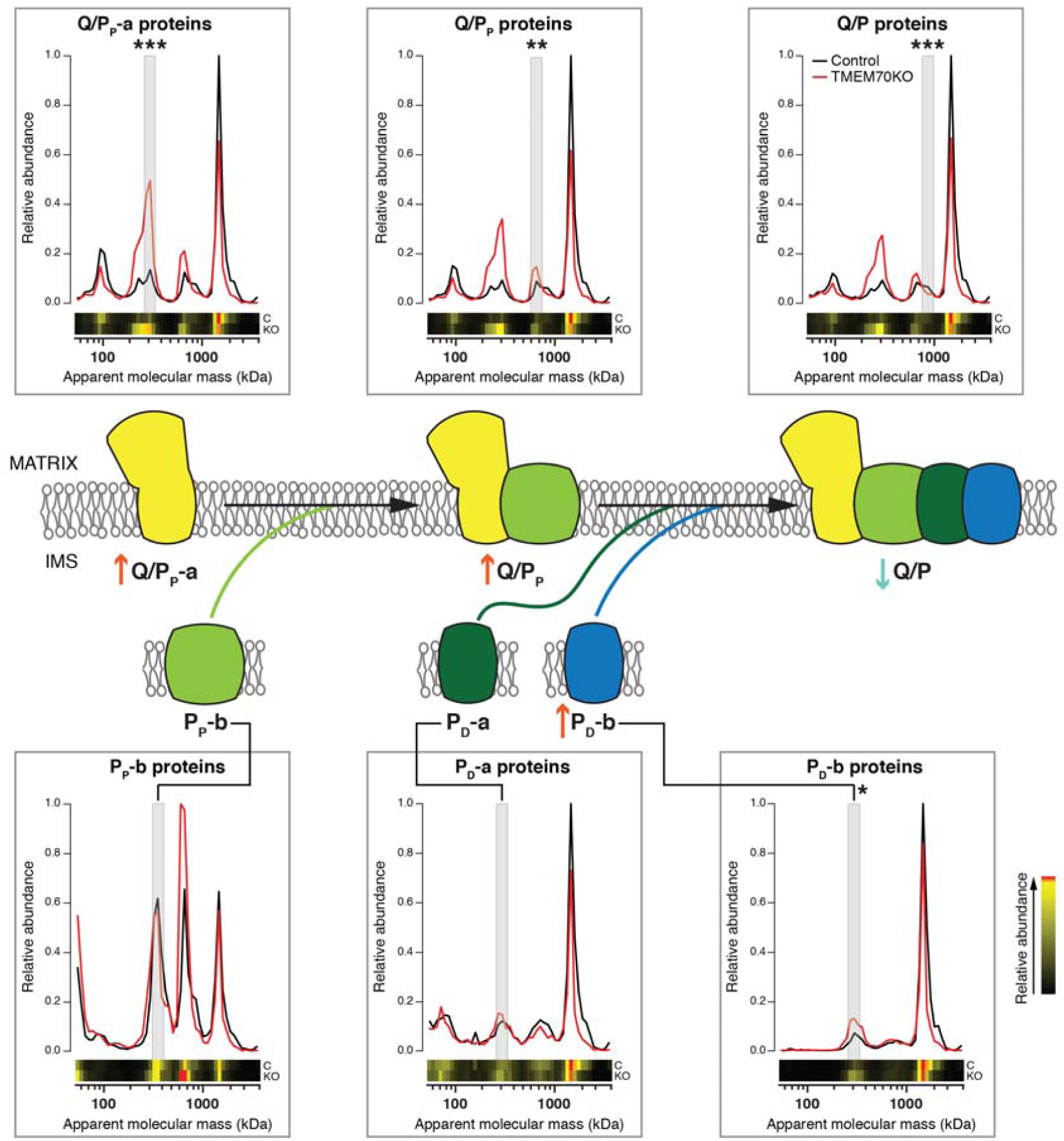
The assembly of complex I is impaired in the absence of TMEM70. Individual plots depict migration profiles of the average of the iBAQ values of the proteins that belong to the stated assembly intermediate (see Methods) in parental HAP1 cells (black line) and TMEM70 KO HAP1 cells (red line). The significant accumulation (red arrow) shown by intermediates Q/P_P_ and P_D_-b together with the significant depletion (blue arrow) of the subsequent product (Q/P) in the TMEM70 knockout cells suggests an impairment in the incorporation of the first two to form the latter. *** *p* < 0.001. ** *p* < 0.01. * *p* < 0.05 based on results depicted in Supplementary Figure S4

### Loss of the interaction between the c-ring and F_1_ module in the *TMEM70* knockout

In order to understand the observed decrease in assembled CV in the absence of TMEM70, we also examined its assembly process in the complexomics data (Supplementary Table S4). We could identify all subunits of CV with two exceptions: the subunit ε (in all samples) and the components of the c-ring octamer (C1 subunits) (in the replicate experiment of the TMEM70 KO cell line). Based on these data, the subunits of CV were distributed over multiple intermediates in the control: the F_1_ soluble and F_1_ anchored intermediates, with subunits α, β, γ and δ in the former, and those plus C1 in the latter (Supplementary Fig. S7a), the F_O_ late assembly intermediate with subunits ATP8, ATP6, DAPIT, 6.8PL, d, F6 and OSCP (Supplementary Fig. S7b) and the F_O_ early assembly intermediate with subunits b, e, f and g (Supplementary Fig. S7c) as well as the fully formed complex V. Subunits belonging to F_1_ show a migration pattern matching previously suggested states^35^, namely, F_1_ soluble (α-β hexamer together with γ, δ and ε monomers), F_1_ attached to the membrane bound c-ring, and F_1_ forming part of the fully assembled complex (Fig. 3a Control, Fig. 3b). With respect to the F_O_ module, our results show a co-migration pattern that hints towards the formation of two intermediates before the fully assembled complex: one formed in an early stage composed of subunits b, e, f and g and one containing these subunits plus the ATP8, ATP6, DAPIT, 6.8PL, d, F6 and OSCP subunits (Supplementary Fig. S7).

**Figure 3.**
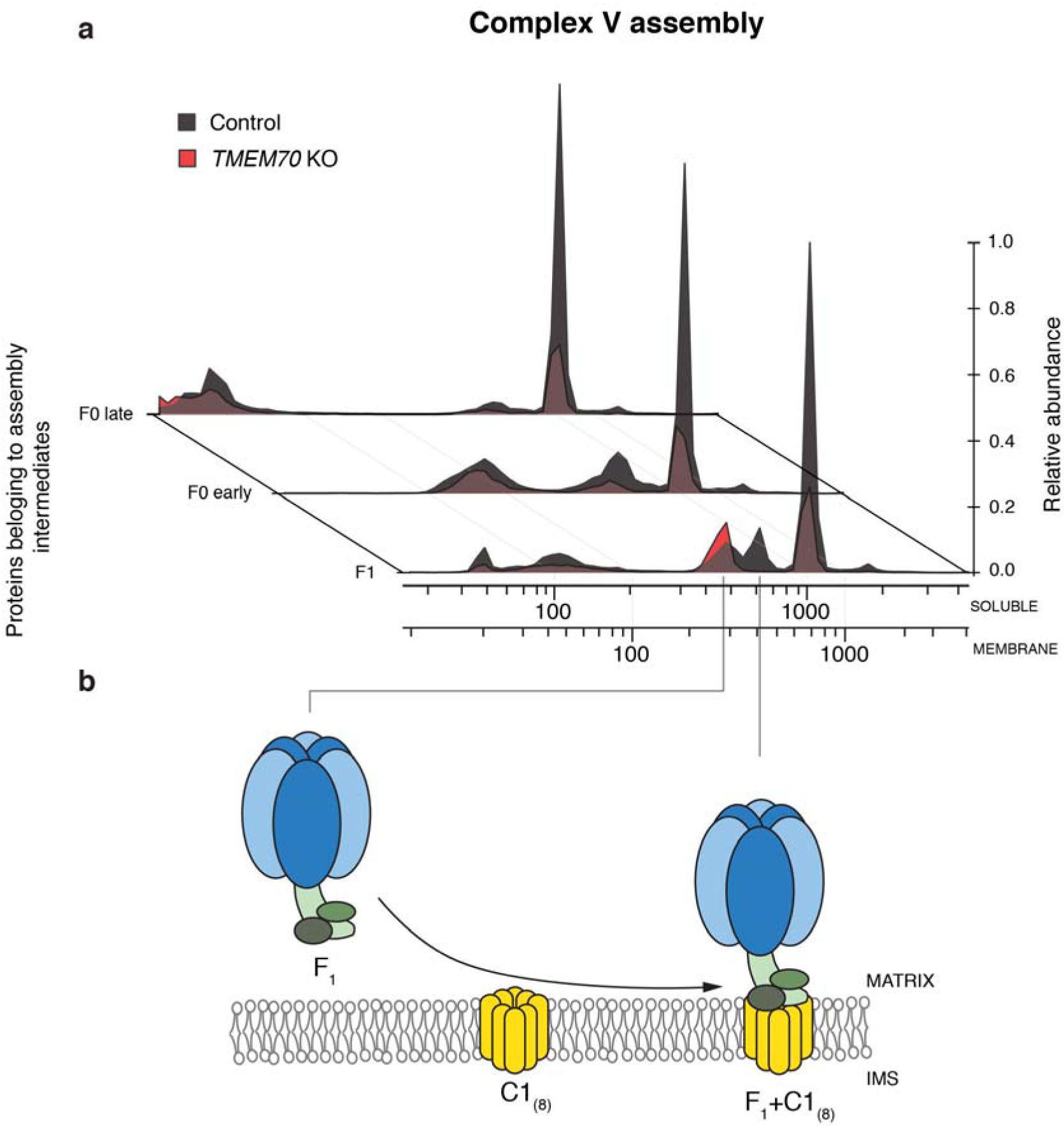
The assembly of complex V misses a F_1_ intermediate in the absence of TMEM70. **a** Migration profiles of complex V intermediates in control HAP1 cells (dark grey) and TMEM70 KO HAP1 cells (red). Intermediates are represented in the z-axis by the average of the iBAQ values of all the subunits detected by mass spectrometry that belong to that intermediate. In absence of TMEM70, F_1_ intermediate accumulation is followed by F_1_-c intermediate depletion whereas the other intermediates do not show specific effects besides an overall reduction of their presence. **b** The F_1_ soluble intermediate is anchored to the inner mitochondrial membrane binding the c-ring octamer

When we explored the assembly of CV in the *TMEM70* knockout cell line in detail, we found that the above-mentioned association of the C1 subunit octamer with the α-β hexamer and the γ, δ and ε subunits that form the F_1_ module, which we observed in control cells, was missing in the absence of TMEM70 (median ratio = 0.04, the small number of subunits does not allow obtaining significant values using non-parametric statistics). Interestingly, the canonical soluble F_1_ intermediate showed a slight accumulation compared to the control (median ratio = 1.27) (Fig. 3a). Consistently, this intermediate was found to be absent in the one-dimensional BN-PAGE immunoblot (Supplementary Fig. S8). With respect to the c-ring components in the complexome analysis by mass spectrometry, we reasoned that the tryptic in-gel digestion may not generate enough detectable peptides of proteins that are small and highly hydrophobic, such as the C1 subunit. To address this issue, we repeated the complexome profiling, but now digesting the samples with chymotrypsin instead of trypsin (Supplementary Table S5). Results revealed that the intermediate in the wild type with an apparent mass of ∼420 kDa indeed represents the F_1_ module associated with the C1 subunit octamer (Fig. 4). Furthermore, the absence of this intermediate in TMEM70 devoid cells was accompanied by the absence of C1 subunit at the same apparent mass, thus confirming the interaction of the soluble F_1_ with the membrane subunit C1 (Fig. 4).

**Figure 4.**
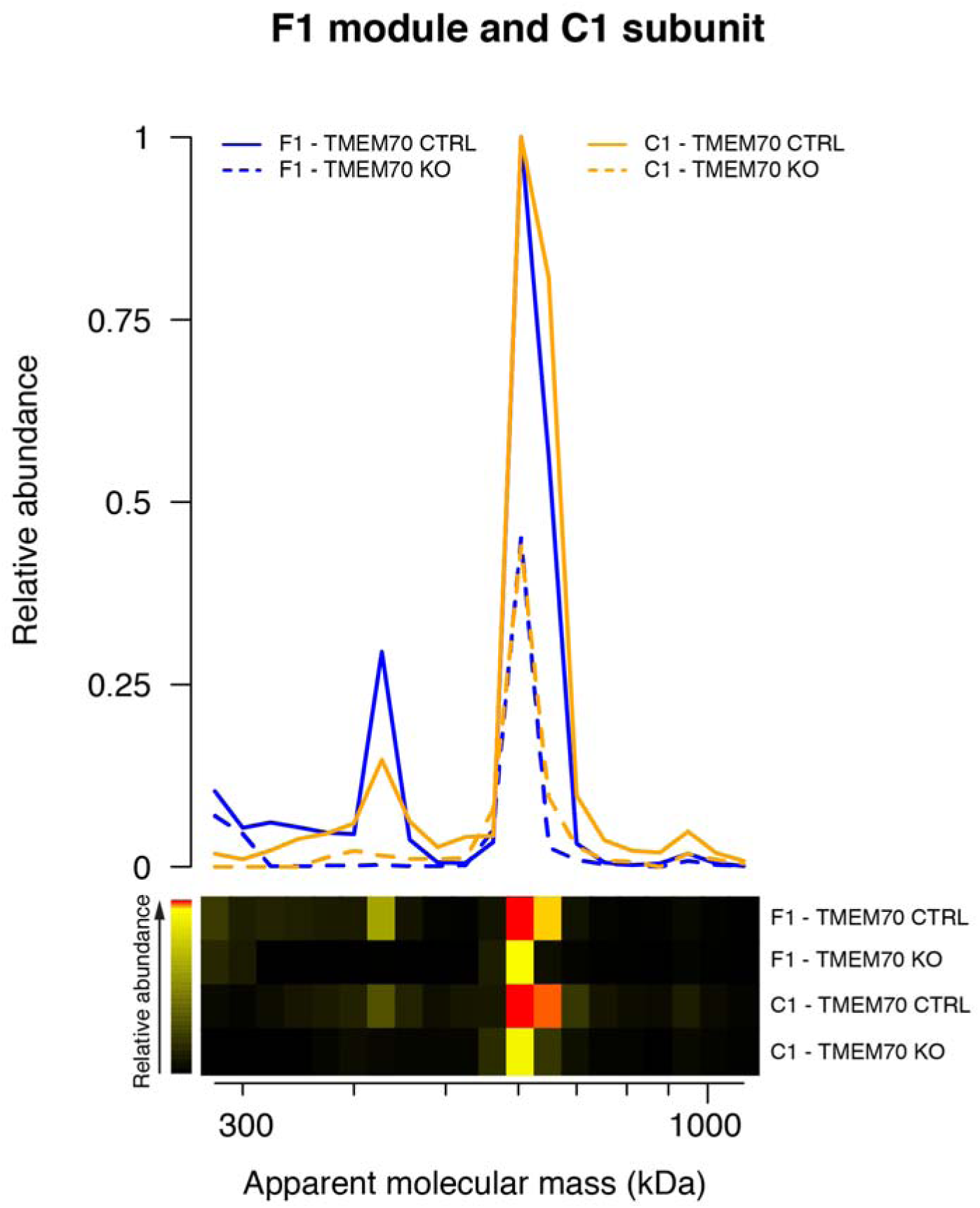
ATP synthase F_O_ complex subunit C1 interaction with F_1_ is missing in the absence of TMEM70. Complexome profiling results after chymotrypsin digestion showing the average of detected F_1_ subunits and C1 protein in control and TMEM70 KO HAP1 cells show a missing intermediate in the latter formed by both groups of proteins (F_1_ proteins and an octamer of C1 subunits).

When focusing on the subunits belonging to the F_O_ early assembly intermediate and/or the F_O_ late assembly intermediate, we observed that the differences between the *TMEM70* knockout and the control are based on an overall reduction (Fig. 3a). Together, these results suggest a role of TMEM70 in the membrane stability of the C1 subunit octamer or its interaction with the F_1_ module.

### Mitochondrial translation is not disturbed by the lack of TMEM70

After confirming the involvement of TMEM70 in complex I and V that was indicated by the BioID results (Table 1), we proceeded with the third complex from the BioID analysis, the mitochondrial ribosome, to determine the possible effect of TMEM70 depletion on the assembly of the small subunit of the mitochondrial ribosomes and/or in mitochondrial translation. To do so, we first assessed the assembly of the small subunit of the mitochondrial ribosomes with our complexomics data in both the presence and absence of TMEM70 (Supplementary Table S4). We did not observe any discrepancies in the formation of intermediates throughout the assembly of the mitochondrial ribosome when comparing both conditions (Fig. 5a), except for a slight but significant increase of the final product (*p* = 7.5e-3, Supplementary Fig. S9) in the absence of TMEM70. In order to assess whether such a difference would affect mitochondrial translation, we labelled newly translated mitochondrial products with ^35^S-methionine^36^ comparing newly synthesized products between control and *TMEM70* KO conditions. We collected cells 5 minutes, 15 minutes, 30 minutes and 60 minutes after 1h of permanently inhibiting cytoplasmic translation (Fig. 5b). Results showed no differences between the two conditions affecting the mitochondrially translated proteins of complexes III and IV. However, we did observe a reduction in ATP6 that becomes more pronounced over time, which is consistent with the strong reduction in fully assembled CV.

**Figure 5.**
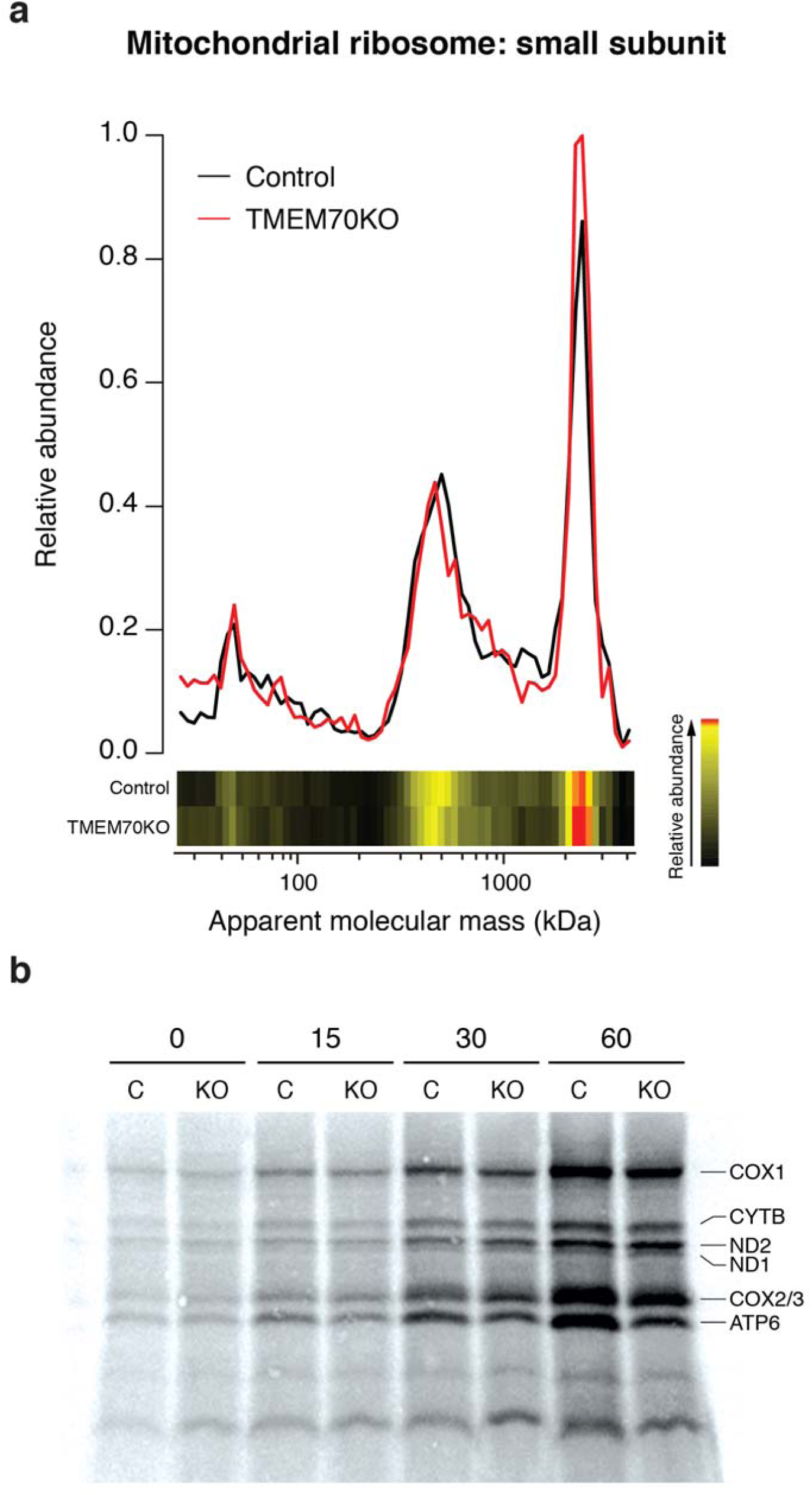
The assembly of the small subunit of the mitochondrial ribosome and the translation of mitochondrial proteins are not affected in the absence of TMEM70 even though there is an increase of the final product. **a** Complexome profile of proteins belonging to the small subunit of the mitochondrial ribosome. Despite being enriched in the BioID dataset, its assembly does not appear to be disturbed in the absence of TMEM70. **b** Pulse labelling of mitochondrial encoded products after 5 min, 15 min, 30 min and 1 h. Newly synthesised products do not seem to show any differences in abundance, with the exception of ATP6 after 15 or more minutes.

The lack of differences in both the assembly process of the ribosomes and the translation efficiency of those suggest no significant role of TMEM70 in their assembly or their normal functioning.

### The mitochondrial protein family TMEM223/TMEM186/TMEM70 co-evolves with OXPHOS

In order examine whether TMEM70 has co-evolved with OXPHOS proteins we determined its phylogenetic distribution. Sequence searches for TMEM70 using sensitive Hidden Markov Models (Methods) revealed that it has two paralogs in human: TMEM223 (E=2.9e-8) and TMEM186 (E=1.2e-28) (Fig. 6a). Results obtained from cellular fractionation (Fig. 6b) and microscopy (Fig. 6c and 6d) indicate a mitochondrial localization of both paralogs. Although the level of sequence identity between the three family members is low (e.g. in human TMEM70-TMEM186: 14%, TMEM70-TMEM223: 9%, TMEM186-TMEM223: 15%), *in silico* predictions indicate a similar asymmetric hairpin topology for all three proteins; a short N-terminal sequence located in the mitochondrial matrix followed by an in/out and an out/in transmembrane helix and a longer C-terminal sequence (Supplementary Fig. S10), consistent with the experimental results for TMEM70^17^. Furthermore, also the *Saccharomyces cerevisiae* protein Mrx15, that is homologous to the TMEM70/186/223 family and within this family most similar to TMEM223 (E=8e-28, 10% identity), has this asymmetric hairpin topology^37^. All three members of the TMEM70/186/223 family are phylogenetically widespread (Fig. 6a), and appear to have been present in the last eukaryotic common ancestor (LECA). We then asked whether any other mitochondrial proteins have a similar phylogenetic distribution as members of the TMEM70/186/223 family. Given the low sequence identity between orthologs in this protein family and between their potential interactors, we used a two-tier approach. First, we employed an in-house orthology database based on pairwise sequence comparisons and containing 52 diverse eukaryotic species to derive phylogenetic profiles of the human proteome, using differential Dollo parsimony that measures the number of independent loss events along an evolutionary tree as an evolutionary distance measure. For TMEM70, the protein with the highest co-evolution signal (NDUFAF1) and four of the ten most co-evolving proteins were CI proteins (Supplementary Table S6). Second, we manually analysed, using the top hits of the first approach and using more sensitive, profile-based analyses (Methods), the co-evolution of TMEM70/186/223 with CI, CV and TMEM14. The latter is, based on the first step of the analysis, a protein family that co-evolves with TMEM70/186/223, and is also located in the inner mitochondrial membrane. Overall, we observe that members of the TMEM70/186/223 protein family only occur in species with OXPHOS, and are absent from species without CI and or CV. Nevertheless, the reverse does not apply as there are taxa, like the Kinetoplastida, that do have CI and CV but do not have detectable homologs of the TMEM70/186/223 family (Supplementary Table S7).

**Figure 6.**
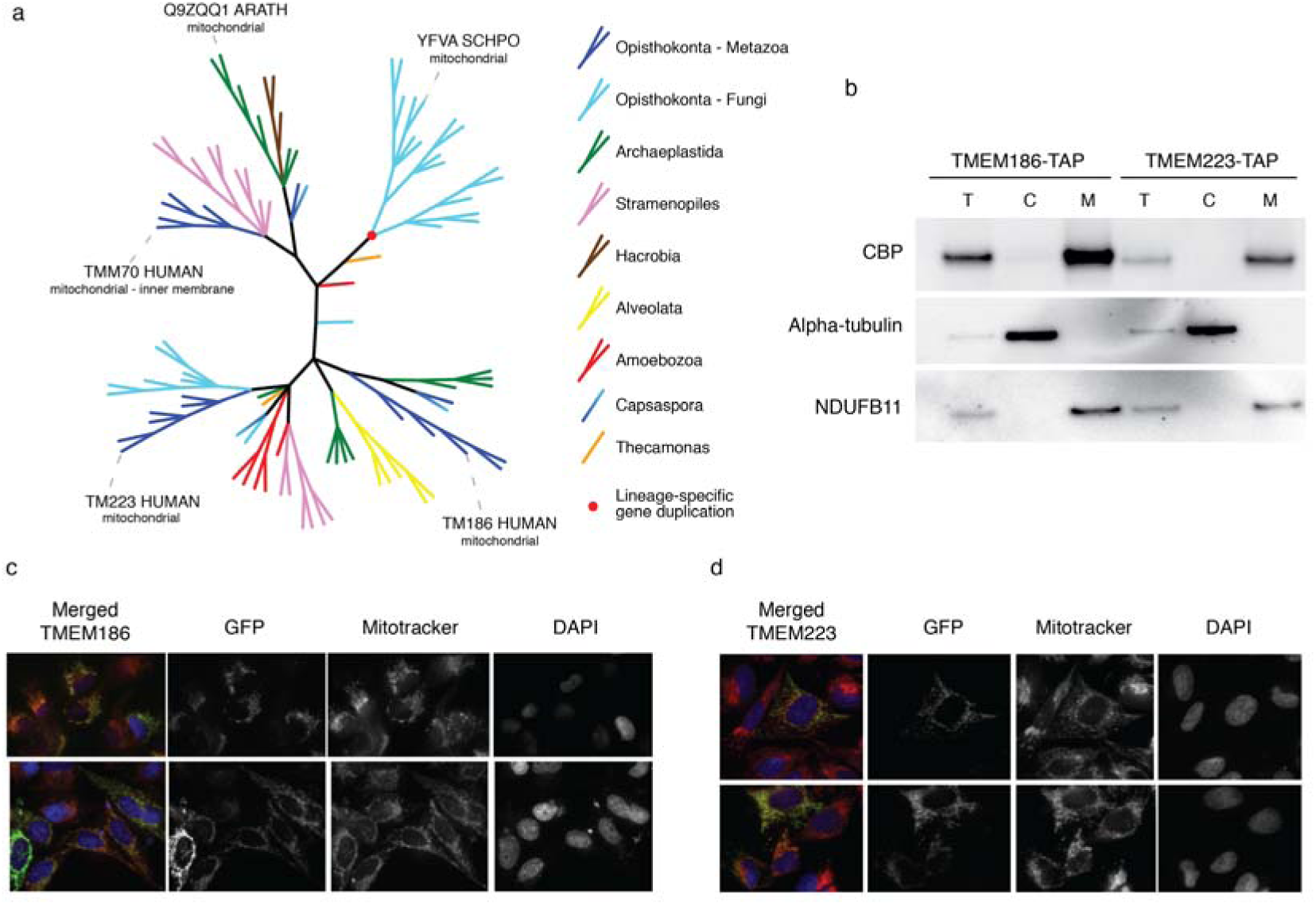
TMEM70 has two paralogs, TMEM186 and TMEM223 that are both located in the mitochondria. Furthermore, for TMEM70 orthologs in *Arabidopsis thaliana* and *Schizosaccharomyces pombe*, there is experimental evidence that localizes them in the mitochondria^38,39^, while also a *S. cerevisiae* homolog of the family, YNR040W/Mrx15 that due to extreme sequence divergence could not be put into the tree, is located in mitochondria^37^. **a** Phylogenetic distribution of TMEM70 homologs. **b** Cellular fractionation of cells expressing TMEM186-TAP and TMEM223-TAP. Immunodetection against NDUFB11, as a marker of the mitochondrial enriched fraction (M), Alpha-tubulin as a marker of the cytoplasm (C), and CBP. (T) stands for total fraction. **c** and **d** Fluorescence microscopy images of cells expressing TMEM186-GFP (**c**) and TMEM223-GFP (**d**) stained with Mitotracker (mitochondrial network) and DAPI (nucleus).

## Discussion

Our results describe a role of TMEM70 in the assembly of CI and CV of the OXPHOS system. This finding is in line with previous studies of patients harbouring mutations in *TMEM70*, that did not always have an isolated CV deficiency but rather combined OXPHOS deficiencies or even an isolated CI deficiency^18,20,22,29^. The conclusions are supported by the results from BioID, complexome profiling analyses and enzyme measurements of the OXPHOS complexes. Although the assembly of both CI and CV are significantly affected in the absence of TMEM70, the effect on CV is more pronounced than on CI, which might explain the preponderance of CV phenotypes relative to CI phenotypes reported for patients with mutations in *TMEM70*.

In order to obtain a comprehensive list of potential interacting candidates that could hint at the function of TMEM70, we began by performing a biotin ligase proximity-dependent assay (BioID) of BirA* tagged TMEM70, which indicated that the protein is in close contact with CI, CV and the small subunit of the mitochondrial ribosome. To determine a potential role of TMEM70 in the assembly of CI and CV in detail, we studied all the proteins of both complexes meticulously by complexome profiling^31,32^. Thus, we were able to detect the majority of the proteins of each complex and of their assembly intermediates. This allowed us to determine the effects that the lack of TMEM70 has on CI and CV assembly. For CI, the Q/P_P_-a and Q/P_P_ intermediates accumulate in the absence of TMEM70, suggesting that the impairment in the pathway comes right after their assembly. In the case of Q/P_P_-a, and following one of the alternative assembly pathways (Supplementary Fig. S6), its accumulation could be caused by a reduction of forming the intermediate P_P_-b/P_D_-a. If we follow the other alternative assembly pathway (Fig. 2), intermediate Q/P_P_ accumulation followed by Q/P depletion, suggests a problem with the incorporation of P_D_-a and P_D_-b intermediates, the latter of which also accumulates in the TMEM70 KO. The interaction between CI and TMEM70 is in line with a previous study by Guarini *et al.*^40^, who showed TMEM70 as a significant interactor of ECSIT and NDUFS5, both proteins belonging to the P_P_-b/P_D_-a intermediate and, in the case of ECSIT, also part of the Q/P_P_ intermediate. The interaction of TMEM70 with proteins of the P_P_ module^40^, the comigration with the distal part (P_D_) of CI^6^, and the accumulation of Q/P_P_-a, Q/P_P_ and P_D_-b intermediates together with the depletion of P_P_-b/P_D_-a and Q/P intermediates, suggests that TMEM70 is involved in the assembly of the membrane arm (P-module) of CI by assisting in assembly or the stabilization of the P_D_-a intermediate A role in the assembly or the stability of a membrane intermediate is consistent with the membrane architecture of the protein, consisting of two transmembrane helices followed by a long C-terminal tail^22^.

With respect to CV, we delineate a specific role of TMEM70 in its assembly. When subunits of F_1_ were affected by a mutation^41^ or were knocked down^42^, an accumulation of subunit c was reported. However, such effect was not observed in patients harbouring mutations in TMEM70^41^, which is consistent with our TMEM70 KO data. Moreover, both in the complexomics data and in the two dimensional BN-PAGE we show how, upon lack of TMEM70, the intermediate where F_1_ is bound to the membrane-embedded c-subunits octamer^35^ is completely absent. While these results are consistent with the ones reported by Vrbacký *et al* in *Tmem70* knockout mouse embryos^28^, by detecting the F_1_-c intermediate in the presence of TMEM70 and its disappearance in the absence of this protein, we are able to specifically link TMEM70 to the formation of the F_1_-c assembly intermediate. This absence can be regarded as an immediate consequence of TMEM70’s impairment and, together with the aforementioned accumulation of c subunits when components of the F_1_ module are defective, indicate that TMEM70 stabilizes the c-ring possibly in conjunction with tethering it to the F1 module (see below). Despite the absence of the F_1_-c intermediate, there was still some CV being assembled, although at only 30% compared to the control. One possibility is that in TMEM70 depleted cells the assembly of the complex follows an alternative path that was already described by He *et al*^43^, where F_1_ instead of being attached to the c-ring octamer first, instead attaches to the F_O_ and the peripheral stalk before combining with the c-ring. Nevertheless, in our complexome profiling data we do not detect such an F_1_F_O_ intermediate that would support this alternative path. With respect to the small subunit of the mitochondrial ribosome, although we observed a slight but significant increment of mitochondrial ribosomes upon *TMEM70* knockout, we did not observe any detectable differences in its assembly nor in the efficiency of the translation of the mitochondrial products. Thus, as the synthesis of mitochondrial encoded subunits was not altered in TMEM70 devoid cells, we find no evidence for the hypothesis of nucleoid disruption^44^. The interactions detected by the BioID experiment could be due to the structure of TMEM70 and the addition of the BirA domain to its C-terminus. Such terminus protrudes into the mitochondrial matrix after the two transmembrane domains. This, together with the possibility of mitochondrial ribosomes being in close proximity to the inner membrane during the translation, could be the reason they were tagged.

Through our homology analyses and phylogenetic reconstruction of the co-evolution of TMEM70/186/223 family with CI and CV we could not detect a TMEM70 ortholog in *S. cerevisiae*, which is surprising as *S. cerevisiae* does have CV. Interestingly, it has been described how in this species, F_1_ does not undergo the same assembly steps as in human and does not interact with the c-ring^45^. Furthermore, *S. cerevisiae* lacks CI, fitting our hypothesis of a double role of the protein in human, that is absent from *S. cerevisiae* because one of the target complexes has been lost in evolution (CI), and the manner in which the other target complex is assembled is different than in human. Sensitive homology detection does however reveal the presence of a TMEM70 homolog in *S. cerevisiae*: TMEM223. TMEM70, TMEM223 and the third member of this family that we could detect, TMEM186, appear widespread in eukaryotic evolution and only occur in species with OXPHOS complexes. We performed confocal microscopy for analysis of GFP-tagged TMEM186 and TMEM223. The analysis showed the predominant localization of both proteins in the mitochondria, something that we could confirm by cell fractionation and isolation of mitochondria. Like TMEM70, TMEM186 was also identified to co-migrate with CI assembly intermediates following dynamic complexome profiling^6^. Little is known about this protein; however, it was recently identified as an interaction partner of ECSIT^40^, suggesting a role in CI assembly. Additional studies are required to determine if this protein functions in CI assembly.

It is interesting and unusual that a single protein TMEM70 would be involved in the assembly of two different OXPHOS complexes, CI and CV. A common denominator based on our results is that, in both cases, it appears to recruit proteins to membrane intermediates during the assembly of the enzymes. Such a combination of hydrophobic and hydrophilic interactions would be consistent with the topology of the protein that contains two well conserved transmembrane helices together with a long (∼100 amino acids) tail that protrudes into the matrix. Mrx15, the yeast homolog of TMEM70 and the ortholog of TMEM223, appears to tether the mitochondrial ribosome to the membrane during translation, using the hydrophilic C terminus to interact with the large subunit of the mitochondrial ribosome while the N terminus forms a hairpin in the inner mitochondrial membrane. We hypothesize that TMEM70 has a similar tethering role in the assembly of complexes I and V. With respect to the notable differences observed in the relative amounts of fully assembled complexes I and V the lack of TMEM70 produces and why the former is less affected, we think that the presence of its homolog TMEM186 - which incidentally also seems to comigrate with complex I - might have a similar role thus mitigating the effect of its absence.

## Materials and methods

### Enzyme Measurements

Respiratory chain enzyme analysis in HAP1 were performed as described before^46^. Values are expressed relative to the mitochondrial reference enzyme citrate synthase^47^.

### Cell culture

HAP1 cells were grown in Iscove’s Modified Dulbecco’s Medium (IMDM) in the presence of 10% fetal calf serum and penicillin/ streptomycin.

HEK293 cells were cultured in DMEM (Biowhitaker) supplemented with 10% fetal calf serum (FCS) (v/v) and 1% penicillin/streptomycin (GIBCO). Inducible cell lines were selected on 5 μg/ml blasticidin (Invitrogen) and 200 μg/ml hygromycin (Calbiochem), and for expression of the transgene, 1 μg/ml doxycycline (Sigma Aldrich) was added for 24 hr.

### Knockout

*TMEM70* knockout (HZGHC003615c010) was ordered at Horizon (Austria). A near-haploid human cell line (HAP1) was edited using CRISPR/Cas (Guide RNA sequence: CGGCTGGAGTACGGGGCCTT) resulting in a frameshift mutation of 32bp in exon 1^26^. This results in a 100% knockout. In Table 1 an overview of the knockout cell line is shown.

### Blue-Native, SDS-PAGE Analysis and complex I In Gel Activity Assay

One-dimensional 10% sodium dodecyl sulfate-polyacrylamide gel electrophoresis (SDS-PAGE) and 4%–12% blue-native (BN)-PAGE was performed as described previously^48^. Lanes were loaded with 40 (SDS analysis) or 80 (BN analysis) µg of solubilized mitochondrial protein. For the 1D and 2D BN SDS-PAGE followed by immunoblotting, mitoplasts were solubilized with n-dodecyl-β-D-maltoside, whereas for complexome profiling the solubilization was done with digitonin to preserve supercomplexes. After electrophoresis, gels were processed further for in-gel complex I activity assay, in-gel fluorescence detection, immunoblotting, or two-dimensional 10% SDS-PAGE or blotting, proteins were transferred to a PROTAN nitrocellulose membrane (Schleicher & Schuell).

### Antibodies and ECL detection

Immunodetection was performed by the use of the following primary antibodies: CI-NDUFS3, CII-SDHA, CIII-core 2, CIV-COX4, CV-ATPase-α/β (gift from the Molecular Bioenergetics Group, Medical School, Goethe-University Frankfurt^49^), C3ORF60 (NDUFAF3) (Eurogentec) and V5 (Invitrogen). Goat-anti-rabbit and goat-anti-mouse IRDye CW 680 or IRDye CW 800 were used as secondary antibodies, to detect the proteins using the Odyssey system from LI-CORBiosciences. Secondary detection was performed using peroxidase-conjugated anti-mouse or anti-rabbit IgGs (Life Technologies). Immunoreactive bands were visualized using the enhanced chemiluminescence kit (Thermo Scientific) and detected using the Chemidoc XRS+system (Biorad).

### Pulse labelling of mitochondrial translation products

In vitro labelling of mitochondrial translation products was performed as described previously^36^. First, cells were incubated for 30 minutes with methionine, cysteine and glutamine free DMEM (Gibco) supplemented with glutamax (1Xconc), 10% dialyzed FBS and 1.1mg/l sodium pyruvate. Afterwards, emetine was added to a final concentration of 100µg/ml, after 5 minutes 200µCi/ml ^35^S methionine label was added during 1 hour. Cells were refreshed with 10% FBS DMEM media and harvested after 1h and after 5h of labelling and treated for SDS-PAGE or BN-PAGE.

### Generation of inducible cell lines

*TMEM70* was cloned into pDONR201 as described^50^. The complete open reading frame (without stop codon) was created by PCR according to the manufacturer’s protocol and cloned into pDONR201 by the Gateway BP Clonase II Enzyme Mix. The pDONR201-TMEM70 was recombined with the pDEST5-BirA*-FLAG-C-ter using the LR Clonase II Enzyme Mix (Invitrogen). Flp-In T-Rex HEK293 cells were grown in DMEM supplemented with 10% FBS and 1% pen/strep (100 U/ml). One day before transfection Pen/strep is removed. When reached 60-80% confluence cells were transfected with each of the constructs with pOG44 using Superfect. Cells were refreshed 3h after transfection. After 48h cells were selected by the addition of hygromycin and Blasticidin (Invitrogen). Clones were selected and when reaching 60-80% confluency, protein expression was induced by using 1 μg/ml doxcycline (Sigma Aldrich) for 24h and checked by Western blot analysis.

### Biotin ligase proximity assay (BioID)

Doxycycline 1 μg/ml (Sigma Aldrich) was added to Flp-In T-Rex HEK293 cells to induce the expression of the construct. After 24h the cells were washed with PBS 1X twice and placed in complete medium for 3h. Afterwards, a final concentration of 50 μM of biotin was added to the cells to start the biotinylation process ^51^. After 24h cells were washed, collected and lysed in lysis buffer (50 mM Tris-HCl pH7.4, 500 mM NaCl, 0.4% SDS, 1 mM DTT and 1x Protease inhibitor) with 2% TX100 and then sonicated twice. Pre-chilled 50 mM Tris-HCl pH7.8 was added before the third sonication and then all samples were collected at 16.500g for 10 minutes at 4 °C. Supernatant was added to the pre-equilibrated dynabeads and it was incubated overnight in the rotator. Next day, beads were collected in the magnetic separation stand and the supernatant was removed. Beads were washed 4 times with 4 different washing buffers (Wash buffer 1: 2% SDS. Wash buffer 2: 0.1% (w/v) deoxycholic acid, 1 % TX100, 1 mM EDTA, 500 mM NaCl, 50 mM HEPES pH 7,4. Wash buffer 3: 0.5 % (w/v) deoxycholic acid, 0.5 % (w/v) NP-40 (Igepal), 1 mM EDTA, 250 mM LiCl, 10 mM Tris-HCl pH7.8) for 8 minutes in the rotator. After the washing 50 μl 50mM ABC / 8M Urea was added to the beads and resuspended gently by pipetting, then the sample was snap frozen in LN2 and stored at −80 degrees.

To prepare the beads first we added 1 μl of reduction buffer (10mM DTT) for 30’ at RT, then 1 μl alkylation buffer (50 mM chloroacetamide in 50mM ABC) and incubated for 20’ at room temperature (light protected). Afterwards 1ug of LysC was added and incubated for at least 3 hours at room temperature. Subsequently the sample was diluted in 50mM ABC and 1ul of trypsin was incorporated for digestion overnight at 37 degrees on a thermomixer on continuous agitation at 700 rpm. The supernatant containing the peptides was transferred to a new Eppendorf tube and TFA was added to a final concentration of 2%. All samples were incorporated into STAGE TIPS that were previously washed, once all sample has pass the filter 50 µl of buffer A (0.1% formic acid in HPLC water) was added and centrifuged. Then 40 µl of buffer B (80% ACN and 0.1% formic acid in HPLC water) was added and the eluate was collected in 0.5 ml reaction vials. All samples were concentrated and dried. Afterwards, 25 μl of buffer A was added and sonicated 2’ in water bath before using the detergent removal kit. Measurements were performed by nanoLC 1000 (Thermo Scientific) chromatography coupled online to Q Exactive hybrid quadrupole-Orbitrap mass spectrometer (Thermo Scientific). Chromatography was performed with an Acclaim PepMap 0.3 × 5 mm 5μm 100Å trap column (Thermo scientific) in combination with a 15cm long × 100μm ID fused silica electrospray emitter (New Objective, PicoTip Emitter, FS360-100-8-N-5-C15) packed in-house with ReproSil-Pur C18-AQ 3 μm 140Å resin (Dr. Maisch)^52^. Tryptic peptides were loaded and analysed by liquid chromatography tandem mass spectrometry (LC-MS/MS) as previously explained. Raw data files provided by MaxQuant (version 1.5.0.25; www.maxquant.org) were further analysed taking the log10 value of their iBAQ and comparing experiment values (with doxycycline and biotin) with control ones (without doxycycline and with biotin and vice versa). Proteins with a significant increase in their iBAQ values (Wilcoxon rank-sum test, FDR 0.05) with respect to their own control runs, were considered as potential interacting candidates.

### Complexome profiling

Complexome profiling was performed essentially as described previously^6,32^, except for the chymotrypsin complexomics data, where chymotrypsin was used instead of trypsin and, from the 60 slices, 20 (slices 30 to 50) where further processed.

BN-PAGE gel lanes of mitochondrial enriched fraction of *TMEM70* KO and controls were incubated in fixing solution (50% methanol, 10% acetic acid, 10 mM ammonium acetate [pH 3]) for 60 minutes, washed twice for 30 minutes with ultrapure water, cut in 60 even slices and transferred into a 96-well plate (Millipore MABVN1250) containing 150µl of destaining solution (50% methanol, 50 mM ammonium hydrogen carbonate, AHC). Then all the slices were washed three times for 30 minutes under gentle agitation at room temperature with the same solution (AHC) to remove the excess of dye. Afterwards by centrifugation (600 x g, 3min, RT) the excess of solution was removed and in-gel tryptic digest of the gel slices was performed as described before^6,32^. Gel slices were incubated with 120 µl of 5mM dithiothreitol for 60 minutes which afterwards was removed by centrifugation, and then 120µl of 15mM chloroacetamide were added to each well and also removed after 45 minutes of incubation. After drying the gel pieces for 45 minutes at room temperature the gel slices were rehydratated in 20µl of trypsin solution or chymotrypsin (5 ng/µL in 50 mM AHC and 1 mM CaCl_2_) for 30 min at 4°C. Alternatively, chymotrypsin was used in an independent replicate. After addition of 150µl of 50mM AHC solution the gel pieces were incubated overnight at 37°C. The supernatant, containing the peptides was collected by centrifugation (600 x g, 3min, RT) into a new 96-well plate. The column was washed subsequently with 80% acetonitrile and re-equilibrated with 5% acetonitrile for 5 minutes once by adding elution solution (30% acetonitrile, 3% formic acid) for 20 minutes and transferred to a sterile (unfiltered) 96-well plate. The supernatant was dried using a SpeedVac concentrator, remaining peptides were additionally extracted in 20 µl of 5% acetonitrile/0.5% formic acid.

Tryptic peptides were analysed by liquid chromatography tandem mass spectrometry (LC-MS/MS) in a Q-Exactive Orbitrap Mass Spectrometry System equipped with a nano-flow high-performance liquid chromatography system EASY-nLC 1000 at the front end and the Thermo Scientific Xcalibur 2.2 SP1 Software Package. Peptide separation was performed on a PicoTip emitter column filled with 3 mm C18 beads (Dr Maisch GmbH, Germany) using 30 min linear gradients of 5 to 35% acetonitrile with 0.1% formic acid. The mass spectrometer was operated in a Top 20 dependent, positive ion mode switching automatically between MS and MS/MS. Full scan MS mode (400 to 1400 m/z) was operated at a resolution of 70 000 with automatic gain control (AGC) target of 1 × 10^6^ ions and a maximum ion transfer of 20 ms. Selected ions for MS/MS were analysed using the following parameters: resolution 17500; AGC target of 1 × 10^5^; maximum ion transfer of 50 ms; 4.0 m/z isolation window; for CID a normalized collision energy 30% was used; and dynamic exclusion of 30.0 s. A lock mass ion (m/z=445.12) was used for internal calibration^53^.

All raw files were analyzed by MaxQuant software (version 1.5.0.25; www.maxquant.org). Spectra were searched against the *H. sapiens* NCBI RefSeq database with additional sequences of known contaminants and reverse decoy with a strict FDR of 0.01. Database searches were done with 20 ppm and 0.5 Da mass tolerances for precursor ions and fragmented ions, respectively. Trypsin was selected as the protease with two missed cleavages allowed. Dynamic modifications included N-terminal acetylation and oxidation of methionine. Cysteine carbamidomethylation was set as fixed modification. For protein quantification, unique plus razor peptides were considered.

### Normalization and alignment of complexome profiles

After analysis in Maxquant, the complexome profiles were normalised to correct for varying intensities. Profiles were corrected so that the sum of intensities of proteins annotated as mitochondrial in MitoCarta 2.0^33^ are equal between samples. After normalisation, the profiles were aligned in silico with COPAL^34^ to correct for technical variation caused by shifts in protein migration across the gel. Gaps introduced in the alignment were filled by linear interpolation based on its adjacent values.

### Quantification of fully assembled complexes and intermediates

Normalized iBAQ values were taken from the profiles after aligning them with COPAL^34^ (Supplementary Table S4). In the case of fully assembled complexes, subunits belonging to CI (NDUFB3, NDUFC2, NDUFB1, NDUFA5, NDUFA6, NDUFS4, NDUFB10, NDUFB2, NDUFB5, NDUFS1, NDUFS8, NDUFAB1, NDUFA2, NDUFS2, NDUFS3, NDUFB6, NDUFB4, NDUFB8, NDUFA7, NDUFB7, NDUFV1, NDUFA8, NDUFV3, NDUFA3, NDUFC1, NDUFS5, NDUFA1, NDUFS6, NDUFV2, NDUFS7, NDUFA11, NDUFA9, NDUFB11, NDUFA13, NDUFA12, NDUFB9, ND1, ND2, ND3, ND4, ND5, ND6), CII (SDHA, SDHB, SDHC, SDHD), CIII (CYTB, UQCRQ, UQCR11, UQCRH, UQCRB, UQCRC2, UQCRC1, UQCRFS1, CYC1, UQCR10), CIV (COX6C, COX3, COX2, COX5B, COX4I1, COX6B1, COX7C, COX5A, COX7B, COX7A2, COX8A, COX1), CV (ATP8, ATP6, ATP5G1, ATP5H, ATP5L, ATP5J, ATP5D, ATP5O, ATP5J2, C14orf2, ATP5I, ATP5F1, ATP5C1, USMG5, ATPIF1, ATP5A1, ATP5B) were considered. With respect to CI intermediates, subunits belonging to all the previously described intermediates of CI assembly^6^, namely, Q/P_P_-a (NDUFA5, NDUFS2, NDUFS3, NDUFS7, NDUFS8, NDUFAF4, NDUFAF3, TIMMDC1, ND1, NDUFA3, NDUFA8, NDUFA13), P_P_-b (NDUFA5, NDUFS2, NDUFS3, NDUFS7, NDUFS8, NDUFAF4, NDUFAF3, TIMMDC1, ND1, NDUFA3, NDUFA8, NDUFA13), P_D_-a (NDUFB6, NDUFB5, NDUFB10, NDUFB11, NDUFB1, ND4, FOXRED1, ATP5SL), P_D_-b (NDUFAB1, NDUFB7, NDUFB3, NDUFB8, ND5, NDUFB9, NDUFB2), Q/P_P_ (NDUFA5, NDUFS2, NDUFS3, NDUFS7, NDUFS8, NDUFAF4, NDUFAF3, TIMMDC1, ND1, NDUFA3, NDUFA8, NDUFA13, NDUFAF2, NDUFA9, NDUFA1), P_P_-b/P_D_-a (NDUFS5, NDUFB6, NDUFB5, NDUFB10, NDUFB11, NDUFB1, ND4, FOXRED1, ATP5SL, NDUFB4) and Q/P (NDUFA5, NDUFS2, NDUFS3, NDUFS7, NDUFS8, NDUFAF4, NDUFAF3, TIMMDC1, ND1, NDUFA3, NDUFA8, NDUFA13, NDUFAF2, NDUFA9, NDUFA1, NDUFB6, NDUFB5, NDUFB10, NDUFB11, NDUFB1, ND4, FOXRED1, ATP5SL, NDUFAB1, NDUFB7, NDUFB3, NDUFB8, ND5, NDUFB9, NDUFB2, NDUFS5) were considered. Regarding CV intermediates, subunits belonging to intermediates described in Supplementary Fig. S7 were considered. Those are F_1_ and F_1_-c (ATP5A1, ATP5B, ATP5C1, ATP5D), F_O_ early (ATP5F1, ATP5I, ATP5J2, ATP5L) and F_O_ late (ATP6, ATP8, C14orf2, USMG5, ATP5H, ATP5J, ATP5O, ATP5F1, ATP5I, ATP5J2, ATP5L). For each subunit, we considered and averaged all three values from the peak matching the mass of the complex and its flanking values. Then, we averaged them per condition, finally obtaining one value per subunit. We compared these values between control and KO conditions in a paired manner per subunit using a non-parametric test (Wilcoxon rank-sum test). For intermediates Q/P_P_ and P_P_-b/P_D_-a, given their partial overlap due to their similar mass, we selected subunits that are not shared between them in order to be able to measure their behaviour independently.

### Microscopy

For confocal imaging, HEK293 cells expressing inducible NDUFAF3-GFP were cultured in a Wilco dish (Intracel, Royston, UK), washed with phosphate-buffered saline (PBS), and incubated with 1 μM Mitotracker Red (Invitrogen) for 15 min and with 10 μM Hoechst 3342 (Invitrogen) for 30 min, both at 37°C. Before imaging, the culture medium was replaced by a colorless HEPES-Tris (HT) solution (132 mM NaCl, 4.2 mM KCl, 1 mM CaCl_2_, 1 mM MgCl_2_, 5.5 mM D-glucose, and 10 mM HEPES, pH 7.4) and fluorescence images were taken on a ZEISS LSM510 Meta confocal microscope (Carl Zeiss). Images were acquired at a rate of 10 Hz with the use of a ×63 oil-immersion objective (N.A. 1.4; Carl Zeiss). Zoom factor 2 and pinhole settings were selected for the attainment of an optical section thickness of < 1 μm. Measurements were performed at 20°C in the dark. Confocal images of GFP and MitoTracker Red fluorescence were simultaneously collected with the use of an argon laser (laser power 1%) with a 488 nm dichroic mirror and a 500–530 nm band-pass barrier filter in combination with a helium-neon (HeNe) 1 laser (laser power 43%) with a 543 nm dichroic mirror and a 560 nm long-pass filter. With the multitrack setting used, Hoechst fluorescence was subsequently imaged with the use of a 405 nm diode laser (laser power 10%) and a 420–480 band-pass barrier filter.

### Phylogenetic reconstruction of TMEM70 family

Protein sequences were manually selected from search results obtained with the jackhmmer tool from HMMER^54^ version 3.1b2. We initiated the search with TMEM70, TMEM186 or TMEM223 and searched against the UniProtKB database, iterating until getting back all three human paralogs (6, 6 and 5 iterations, respectively). In none of the cases did we retrieve other human proteins than TMEM70, TMEM186 or TMEM223. In order to obtain as many sequences from different phyla as possible, we repeated the strategy using the PSI-BLAST algorithm from BLAST^55^ searching against the refseq protein database. After selecting the sequences, we aligned them with MAFFT^56^ v7.306 using the L-INS-I algorithm with the “Leave gappy regions” and “Mafft-homologs: ON” parameters set. The obtained alignment was manually refined by deleting large gaps caused by apparent insertions in only few aligned sequences. Finally, in order to unravel the orthologous groups, we reconstructed the phylogenetic tree with PhyML^57^ version 20120412 and its automatic model selection SMS^58^ using Bayesian Information Criterion. The resulting tree was plotted with iTOL^59^ version 3.5.4.

### Co-evolution screen for TMEM70

We employed an in-house orthology database containing 52 diverse eukaryotic species to derive phylogenetic profiles for all human proteins as previously described by van Dam *et al.*^60^. Shortly, we collected high quality proteomes from genome databases and calculated orthologous groups using OrthoMCL (v2.0)^61^. Phylogenetic profiles were computed for each human protein based on the orthologous group they were mapped to. Proteins mapped to the same orthologous group obtained identical profiles. Each profile is a vector of 0’s and 1’s reflecting the presence of an orthologous group member in a particular species. These profiles, and a species tree for all 52 species as created before^60^ were used as input for the Perl script to derive the differential Dollo parsimony score as previously used by Kensche *et al.*^62^.

## Acknowledgements

Funded by the European Commission (FP7-PEOPLE-ITN. GA. 317433). U.B. and S.G.-C. were supported by a grant from the Excellence Initiative of the German Federal and State Governments (EXC 115). M.A.H., U.B. and J.S. were supported by a TOP grant from the Netherlands Organization for Health Research and Development (no. 91217009). Part of this work was financed by a grant obtained from the United Mitochondrial Disease Foundation (UMDF).

## Author contributions

L.S.-C. and L.G.J.N. conceived the study and designed the research. L.S.-C., M.B. and F.B. performed the experiments and generated the data with help from S.G.-C. and R.R. J.S. aligned the complexomics data. D.M.E. analysed the data. D.M.E. and M.A.H. performed the phylogenetic reconstruction and T.J.P.D. and M.A.H., performed the coevolution analysis. L.S.-C., D.M.E., M.A.H. and L.G.J.N. wrote the manuscript and S.G.-C., J.S., T.J.P.D. R.R. and U.B. discussed the results and helped with the manuscript.

## Competing interests

The authors declare no competing interests.

## References

1 Alston, C. L., Rocha, M. C., Lax, N. Z., Turnbull, D. M. & Taylor, R. W. The genetics and pathology of mitochondrial disease. J Pathol 241, 236-250, doi:10.1002/path.4809 (2017).

2 Mitchell, P. Coupling of phosphorylation to electron and hydrogen transfer by a chemiosmotic type of mechanism. Nature 191, 144-148 (1961).

3 Zhu, J., Vinothkumar, K. R. & Hirst, J. Structure of mammalian respiratory complex I. Nature 536, 354-358, doi:10.1038/nature19095 (2016).

4 Brandt, U. Energy converting NADH:quinone oxidoreductase (complex I). Annu Rev Biochem 75, 69-92, doi:10.1146/annurev.biochem.75.103004.142539 (2006).

5 Sanchez-Caballero, L., Guerrero-Castillo, S. & Nijtmans, L. Unraveling the complexity of mitochondrial complex I assembly: A dynamic process. Biochim Biophys Acta 1857, 980-990, doi:10.1016/j.bbabio.2016.03.031 (2016).

6 Guerrero-Castillo, S. et al. The Assembly Pathway of Mitochondrial Respiratory Chain Complex I. Cell Metab 25, 128-139, doi:10.1016/j.cmet.2016.09.002 (2017).

7 Pedersen, P. L. & Amzel, L. M. ATP synthases. Structure, reaction center, mechanism, and regulation of one of nature’s most unique machines. J Biol Chem 268, 9937-9940 (1993).

8 Walker, J. E. The ATP synthase: the understood, the uncertain and the unknown. Biochem Soc Trans 41, 1-16, doi:10.1042/BST20110773 (2013).

9 Ghezzi, D. & Zeviani, M. Human diseases associated with defects in assembly of OXPHOS complexes. Essays Biochem 62, 271-286, doi:10.1042/EBC20170099 (2018).

10 Elurbe, D. M. & Huynen, M. A. The origin of the supernumerary subunits and assembly factors of complex I: A treasure trove of pathway evolution. Biochim Biophys Acta 1857, 971-979, doi:10.1016/j.bbabio.2016.03.027 (2016).

11 Huynen, M. A., de Hollander, M. & Szklarczyk, R. Mitochondrial proteome evolution and genetic disease. Biochim Biophys Acta 1792, 1122-1129, doi:10.1016/j.bbadis.2009.03.005 (2009).

12 Bych, K. et al. The iron-sulphur protein Ind1 is required for effective complex I assembly. EMBO J 27, 1736-1746, doi:10.1038/emboj.2008.98 (2008).

13 Pickova, A., Paul, J., Petruzzella, V. & Houstek, J. Differential expression of ATPAF1 and ATPAF2 genes encoding F(1)-ATPase assembly proteins in mouse tissues. FEBS Lett 551, 42-46 (2003).

14 Cizkova, A. et al. TMEM70 mutations cause isolated ATP synthase deficiency and neonatal mitochondrial encephalocardiomyopathy. Nat Genet 40, 1288-1290, doi:10.1038/ng.246 (2008).

15 Calvo, S. et al. Systematic identification of human mitochondrial disease genes through integrative genomics. Nat Genet 38, 576-582, doi:10.1038/ng1776 (2006).

16 Hejzlarova, K. et al. Expression and processing of the TMEM70 protein. Biochim Biophys Acta 1807, 144-149, doi:10.1016/j.bbabio.2010.10.005 (2011).

17 Kratochvilova, H. et al. Mitochondrial membrane assembly of TMEM70 protein. Mitochondrion 15, 1-9, doi:10.1016/j.mito.2014.02.010 (2014).

18 Wortmann, S. B. et al. Biochemical and genetic analysis of 3-methylglutaconic aciduria type IV: a diagnostic strategy. Brain 132, 136-146, doi:10.1093/brain/awn296 (2009).

19 Catteruccia, M. et al. Persistent pulmonary arterial hypertension in the newborn (PPHN): a frequent manifestation of TMEM70 defective patients. Mol Genet Metab 111, 353-359, doi:10.1016/j.ymgme.2014.01.001 (2014).

20 Diodato, D. et al. Common and Novel TMEM70 Mutations in a Cohort of Italian Patients with Mitochondrial Encephalocardiomyopathy. JIMD Rep 15, 71-78, doi:10.1007/8904_2014_300 (2015).

21 Honzik, T. et al. Mitochondrial encephalocardio-myopathy with early neonatal onset due to TMEM70 mutation. Arch Dis Child 95, 296-301, doi:10.1136/adc.2009.168096 (2010).

22 Jonckheere, A. I. et al. Restoration of complex V deficiency caused by a novel deletion in the human TMEM70 gene normalizes mitochondrial morphology. Mitochondrion 11, 954-963, doi:10.1016/j.mito.2011.08.012 (2011).

23 Magner, M. et al. TMEM70 deficiency: long-term outcome of 48 patients. J Inherit Metab Dis 38, 417-426, doi:10.1007/s10545-014-9774-8 (2015).

24 Shchelochkov, O. A. et al. Milder clinical course of Type IV 3-methylglutaconic aciduria due to a novel mutation in TMEM70. Mol Genet Metab 101, 282-285, doi:10.1016/j.ymgme.2010.07.012 (2010).

25 Spiegel, R. et al. TMEM70 mutations are a common cause of nuclear encoded ATP synthase assembly defect: further delineation of a new syndrome. J Med Genet 48, 177-182, doi:10.1136/jmg.2010.084608 (2011).

26 Torraco, A. et al. TMEM70: a mutational hot spot in nuclear ATP synthase deficiency with a pivotal role in complex V biogenesis. Neurogenetics 13, 375-386, doi:10.1007/s10048-012-0343-8 (2012).

27 Houstek, J., Kmoch, S. & Zeman, J. TMEM70 protein - a novel ancillary factor of mammalian ATP synthase. Biochim Biophys Acta 1787, 529-532, doi:10.1016/j.bbabio.2008.11.013 (2009).

28 Vrbacky, M. et al. Knockout of Tmem70 alters biogenesis of ATP synthase and leads to embryonal lethality in mice. Hum Mol Genet 25, 4674-4685, doi:10.1093/hmg/ddw295 (2016).

29 Braczynski, A. K. et al. ATP synthase deficiency due to TMEM70 mutation leads to ultrastructural mitochondrial degeneration and is amenable to treatment. BioMed research international 2015, 462592, doi:10.1155/2015/462592 (2015).

30 Huang da, W., Sherman, B. T. & Lempicki, R. A. Systematic and integrative analysis of large gene lists using DAVID bioinformatics resources. Nat Protoc 4, 44-57, doi:10.1038/nprot.2008.211 (2009).

31 Wessels, H. J. et al. LC-MS/MS as an alternative for SDS-PAGE in blue native analysis of protein complexes. Proteomics 9, 4221-4228 (2009).

32 Heide, H. et al. Complexome profiling identifies TMEM126B as a component of the mitochondrial complex I assembly complex. Cell Metab 16, 538-549, doi:10.1016/j.cmet.2012.08.009 (2012).

33 Calvo, S. E., Clauser, K. R. & Mootha, V. K. MitoCarta2.0: an updated inventory of mammalian mitochondrial proteins. Nucleic Acids Res 44, D1251-1257, doi:10.1093/nar/gkv1003 (2016).

34 Van Strien, J. et al. COmplexome Profiling ALignment (COPAL) reveals remodeling of mitochondrial protein complexes in Barth syndrome. Bioinformatics, doi:10.1093/bioinformatics/btz025 (2019).

35 Nijtmans, L. G., Klement, P., Houstek, J. & van den Bogert, C. Assembly of mitochondrial ATP synthase in cultured human cells: implications for mitochondrial diseases. Biochim Biophys Acta 1272, 190-198 (1995).

36 Boulet, L., Karpati, G. & Shoubridge, E. A. Distribution and threshold expression of the tRNA(Lys) mutation in skeletal muscle of patients with myoclonic epilepsy and ragged-red fibers (MERRF). Am J Hum Genet 51, 1187-1200 (1992).

37 Moller-Hergt, B. V., Carlstrom, A., Stephan, K., Imhof, A. & Ott, M. The ribosome receptors Mrx15 and Mba1 jointly organize cotranslational insertion and protein biogenesis in mitochondria. Mol Biol Cell 29, 2386-2396, doi:10.1091/mbc.E18-04-0227 (2018).

38 Nikolovski, N. et al. Putative glycosyltransferases and other plant Golgi apparatus proteins are revealed by LOPIT proteomics. Plant Physiol 160, 1037-1051, doi:10.1104/pp.112.204263 (2012).

39 Matsuyama, A. et al. ORFeome cloning and global analysis of protein localization in the fission yeast Schizosaccharomyces pombe. Nat Biotechnol 24, 841-847, doi:10.1038/nbt1222 (2006).

40 Guarani, V. et al. TIMMDC1/C3orf1 functions as a membrane-embedded mitochondrial complex I assembly factor through association with the MCIA complex. Molecular and cellular biology 34, 847-861, doi:10.1128/MCB.01551-13 (2014).

41 Mayr, J. A. et al. Mitochondrial ATP synthase deficiency due to a mutation in the ATP5E gene for the F1 epsilon subunit. Hum Mol Genet 19, 3430-3439, doi:10.1093/hmg/ddq254 (2010).

42 Pecina, P. et al. Role of the mitochondrial ATP synthase central stalk subunits gamma and delta in the activity and assembly of the mammalian enzyme. Biochim Biophys Acta Bioenerg 1859, 374-381, doi:10.1016/j.bbabio.2018.02.007 (2018).

43 He, J. et al. Assembly of the membrane domain of ATP synthase in human mitochondria. Proc Natl Acad Sci U S A 115, 2988-2993, doi:10.1073/pnas.1722086115 (2018).

44 Cameron, J. M. et al. Complex V TMEM70 deficiency results in mitochondrial nucleoid disorganization. Mitochondrion 11, 191-199, doi:10.1016/j.mito.2010.09.008 (2011).

45 Song, J., Pfanner, N. & Becker, T. Assembling the mitochondrial ATP synthase. Proc Natl Acad Sci U S A 115, 2850-2852, doi:10.1073/pnas.1801697115 (2018).

46 Janssen, A. J. et al. Spectrophotometric assay for complex I of the respiratory chain in tissue samples and cultured fibroblasts. Clin Chem 53, 729-734, doi:10.1373/clinchem.2006.078873 (2007).

47 Srere, P. A. The citrate cleavage enzyme. I. Distribution and purification. J Biol Chem 234, 2544-2547 (1959).

48 Calvaruso, M. A., Smeitink, J. & Nijtmans, L. Electrophoresis techniques to investigate defects in oxidative phosphorylation. Methods 46, 281-287, doi:10.1016/j.ymeth.2008.09.023 (2008).

49 Wittig, I., Velours, J., Stuart, R. & Schagger, H. Characterization of domain interfaces in monomeric and dimeric ATP synthase. Mol Cell Proteomics 7, 995-1004, doi:10.1074/mcp.M700465-MCP200 (2008).

50 Vogel, R. O. et al. Cytosolic signaling protein Ecsit also localizes to mitochondria where it interacts with chaperone NDUFAF1 and functions in complex I assembly. Genes Dev 21, 615-624, doi:10.1101/gad.408407 (2007).

51 Firat-Karalar, E. N. & Stearns, T. Probing mammalian centrosome structure using BioID proximity-dependent biotinylation. Methods Cell Biol 129, 153-170, doi:10.1016/bs.mcb.2015.03.016 (2015).

52 Ishihama, Y., Rappsilber, J., Andersen, J. S. & Mann, M. Microcolumns with self-assembled particle frits for proteomics. J Chromatogr A 979, 233-239 (2002).

53 Olsen, J. V. et al. Parts per million mass accuracy on an Orbitrap mass spectrometer via lock mass injection into a C-trap. Mol Cell Proteomics 4, 2010-2021, doi:10.1074/mcp.T500030-MCP200 (2005).

54 Finn, R. D. et al. HMMER web server: 2015 update. Nucleic Acids Res 43, W30-38, doi:10.1093/nar/gkv397 (2015).

55 Altschul, S. F. et al. Gapped BLAST and PSI-BLAST: a new generation of protein database search programs. Nucleic Acids Res 25, 3389-3402 (1997).

56 Katoh, K. & Standley, D. M. MAFFT multiple sequence alignment software version 7: improvements in performance and usability. Mol Biol Evol 30, 772-780, doi:10.1093/molbev/mst010 (2013).

57 Guindon, S. et al. New algorithms and methods to estimate maximum-likelihood phylogenies: assessing the performance of PhyML 3.0. Syst Biol 59, 307-321, doi:10.1093/sysbio/syq010 (2010).

58 Lefort, V., Longueville, J. E. & Gascuel, O. SMS: Smart Model Selection in PhyML. Mol Biol Evol 34, 2422-2424, doi:10.1093/molbev/msx149 (2017).

59 Letunic, I. & Bork, P. Interactive tree of life (iTOL) v3: an online tool for the display and annotation of phylogenetic and other trees. Nucleic Acids Res 44, W242-245, doi:10.1093/nar/gkw290 (2016).

60 van Dam, T. J. et al. Evolution of modular intraflagellar transport from a coatomer-like progenitor. Proc. Natl. Acad. Sci. U. S. A. 110, 6943-6948, doi:10.1073/pnas.1221011110 (2013).

61 Li, L., Stoeckert, C. J., Jr. & Roos, D. S. OrthoMCL: identification of ortholog groups for eukaryotic genomes. Genome Res. 13, 2178-2189, doi:10.1101/gr.1224503 (2003).

62 Kensche, P. R., van Noort, V., Dutilh, B. E. & Huynen, M. A. Practical and theoretical advances in predicting the function of a protein by its phylogenetic distribution. J R Soc Interface 5, 151-170, doi:10.1098/rsif.2007.1047 (2008).

