## Supplemental Figures for "A dual function of TMEM70 in OXPHOS: assembly of complexes I and V"

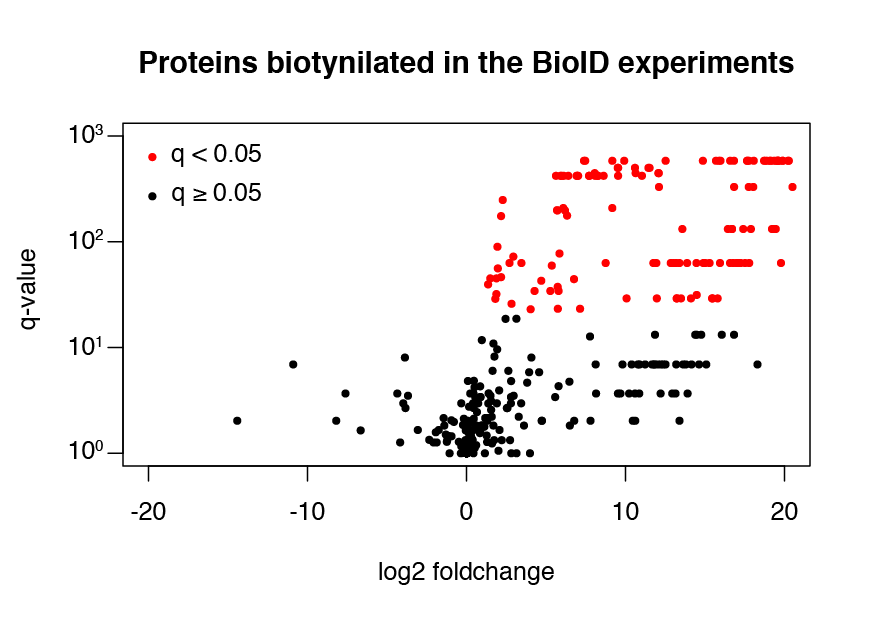

Figure S1. Scatterplot depicting the results of the BioID analysis. Proteins considered positive are depicted in red (q-value < 0.05)

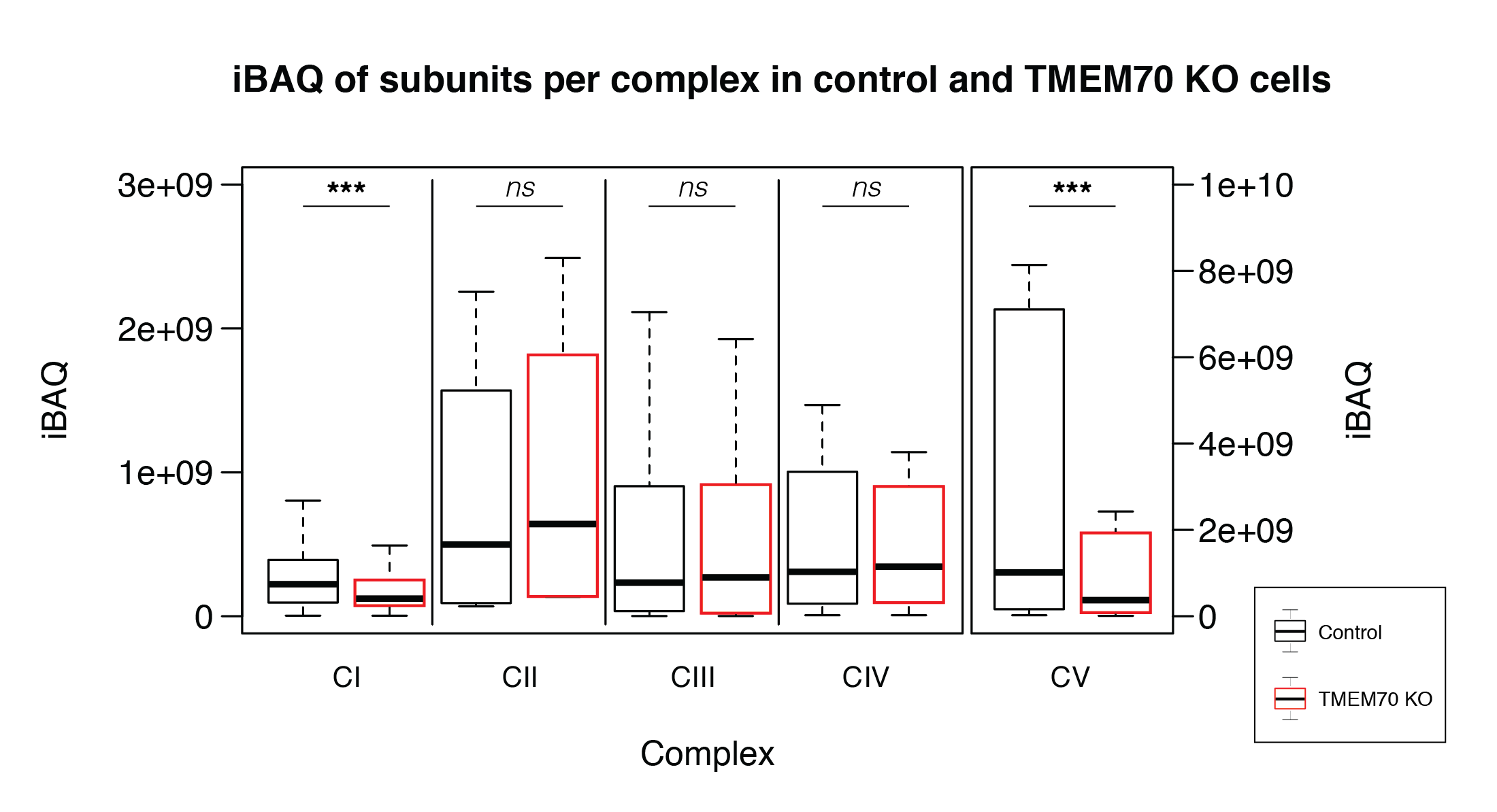

Figure S2. Boxplots depicting the distribution of the iBAQ values of the subunits belonging to each OXPHOS complex. *** *p* < 0.001. n.s. not significant

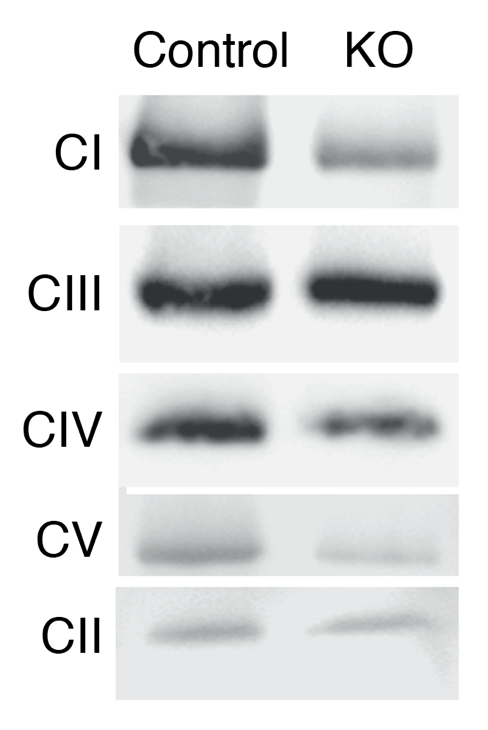

Figure S3. BN-PAGE/immunoblotting analysis of CI (NDUFS3), CII (SDHA), CIII (UQCRC2), CIV (COX4) and CV (α subunit) reveals deficiency of CI and CV in TMEM70 depleted HAP1 cells (KO) compared with its parental HAP1 (Control). Levels of complex II (CII, SDHA) serve as loading control.

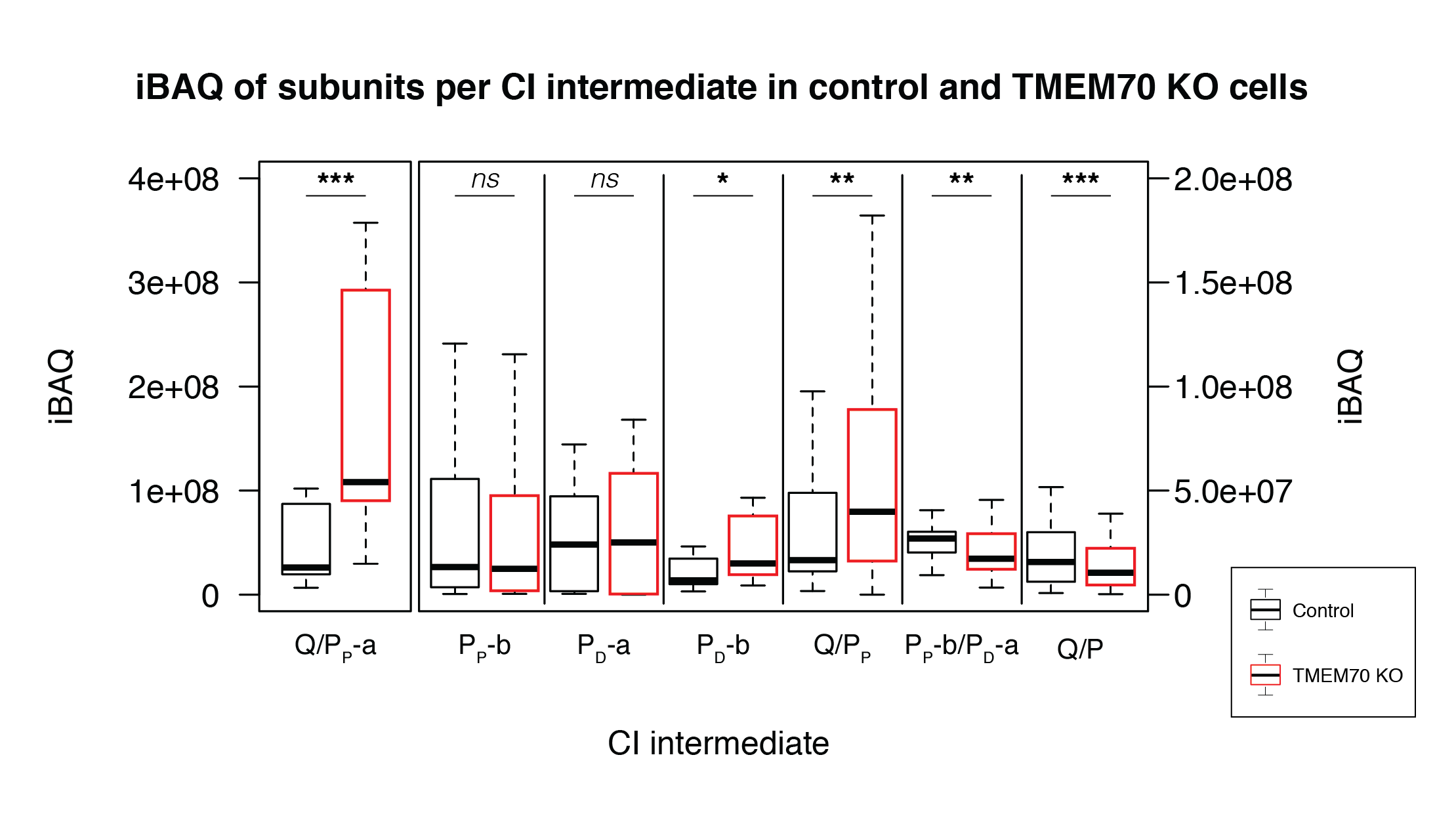

Figure S4. Boxplots depicting the distribution of the iBAQ values of the subunits belonging to each CI assembly intermediate. *** *p* < 0.001. ** *p* < 0.01. * p < 0.05. n.s. not significant

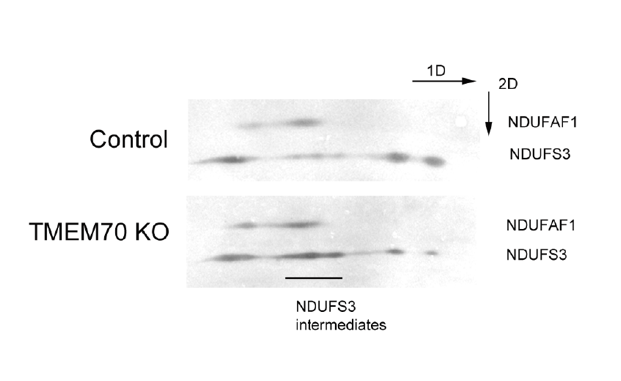

Figure S5. Two-dimensional Blue Native PAGE analysis of HAP1 control cells (Control) and HAP1 TMEM70 depleted cells (TMEM70 KO). (Mitoplasts were lysed with n-dodecyl- β-D-maltoside. Blots were stained with antibodies directed against representative subunits of different CI intermediates: NDUFS3 (Q-module) and NDUFAF1 (P_P_-b) to detect Q/P_P_ intermediates.

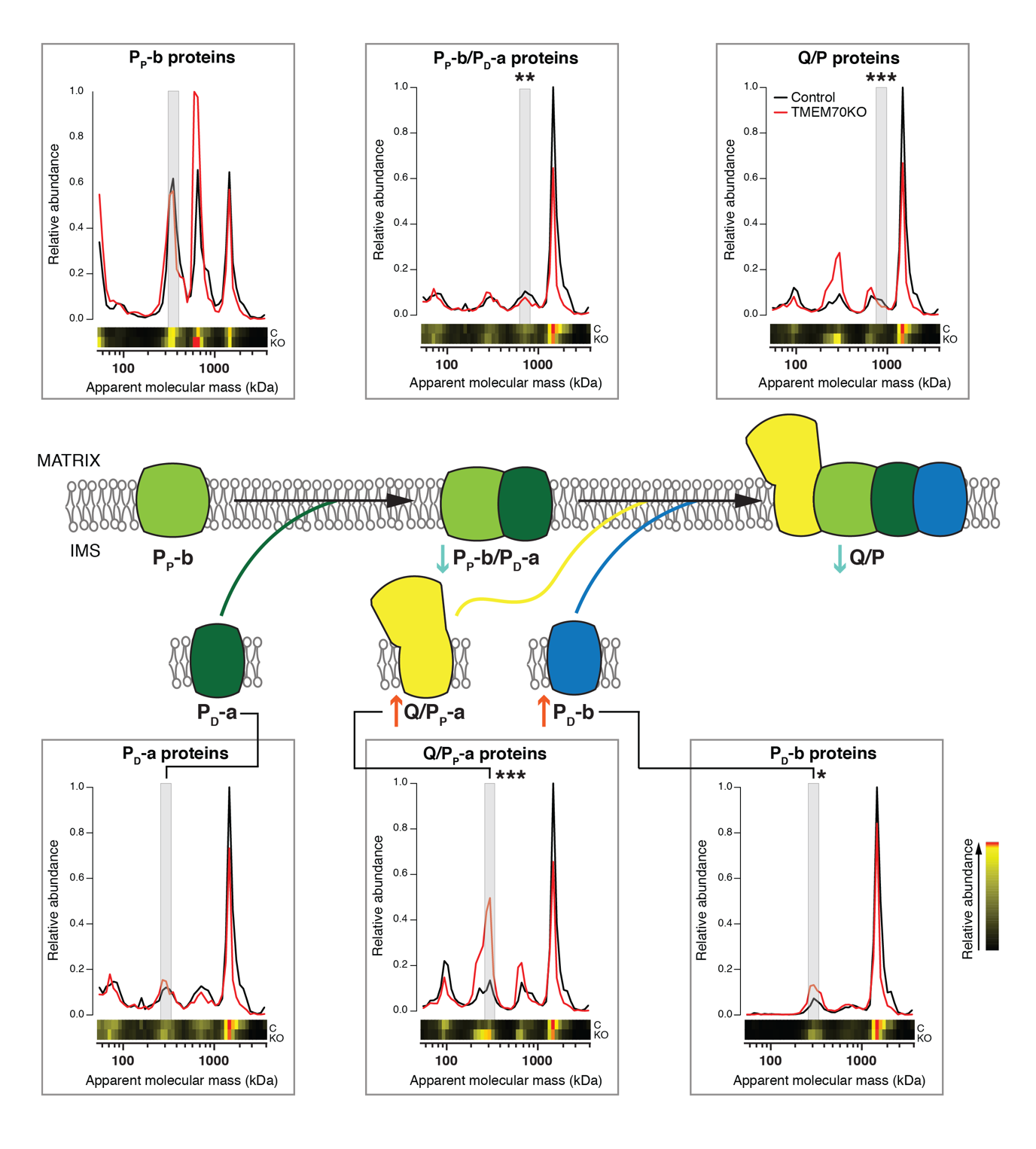

Figure S6. The alternative assembly pathway of complex I also shows depletion of intermediates after the addition of P_D_-a in the absence of TMEM70**.** Individual plots depict migration profiles of the average of the iBAQ values of the proteins that belong to the stated assembly intermediate (see Methods) in parental HAP1 cells (black line) and TMEM70 KO HAP1 cells (red line). The significant accumulation (red arrow) shown by intermediates Q/P_P_-a and P_D_-b is also consistent with the significant depletion (blue arrow) of intermediates P_P_-b/P_D_-a and Q/P in the TMEM70 knockout cells. *** *p* < 0.001. ** *p* < 0.01. * *p* < 0.05 based on results depicted in Supplementary Figure S4

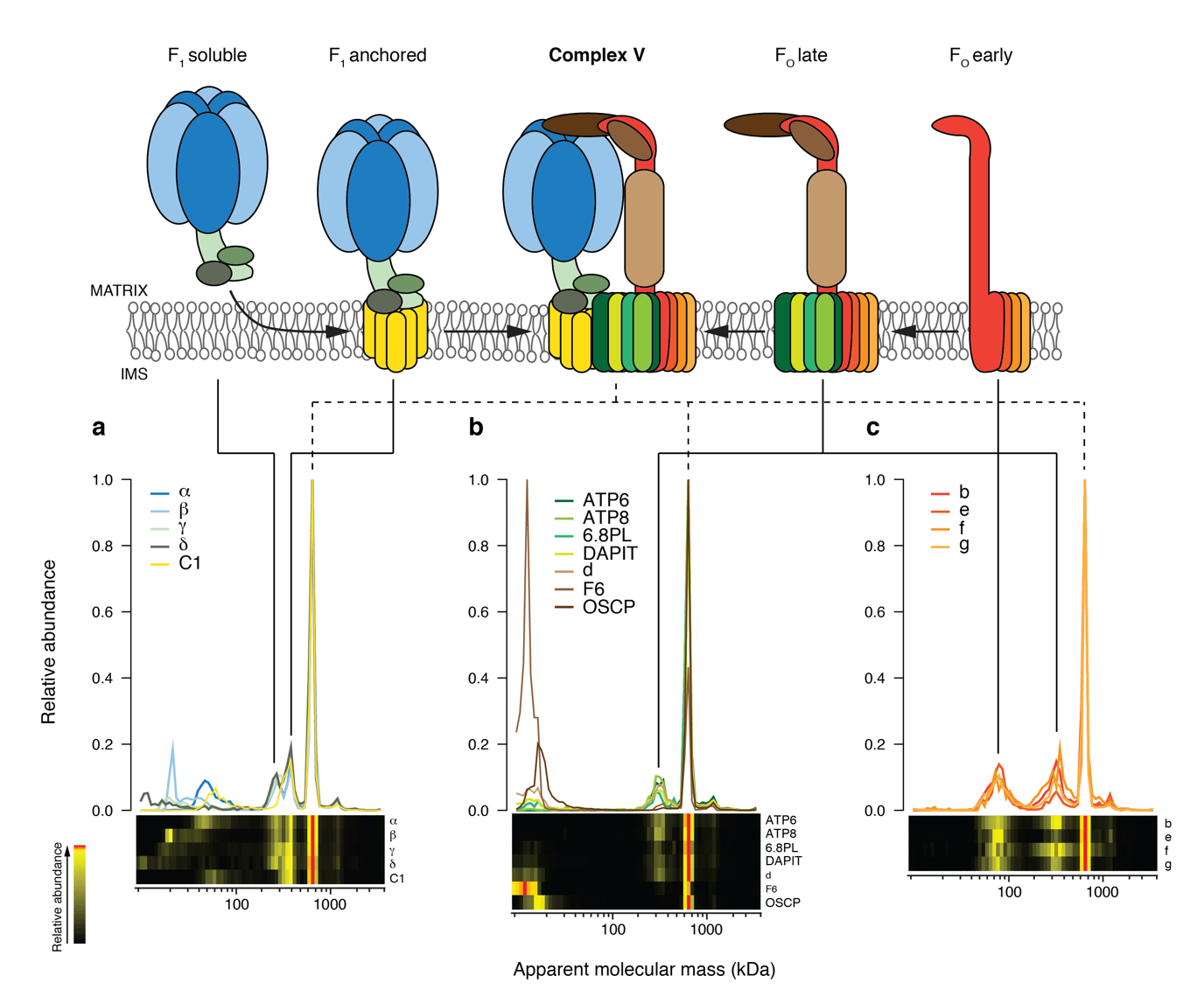

Figure S7. Assembly of complex V based on the migration profile of the proteins that belong to the enzyme obtained by complexomics on HAP1 control cells. **a** F_1_ module subunits assemble and are afterwards anchored to the membrane with subunit C1. **b** Subunits of the F_O_ module that are incorporated in a later stage of its assembly. **c** Subunits of the F_O_ module that form the first intermediate of such module

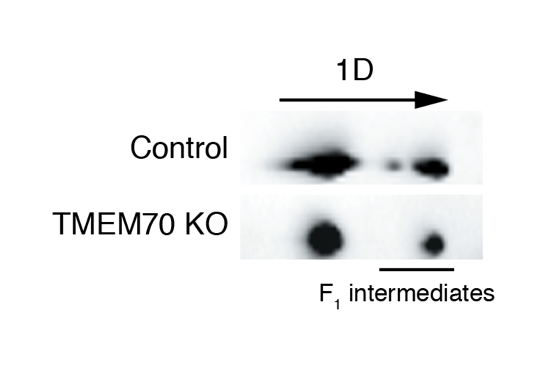

Figure S8. Two-dimensional Blue Native PAGE analysis of HAP1 control cells (Control) and HAP1 TMEM70 depleted cells (TMEM70 KO). (Mitoplasts were lysed with n-dodecyl- β-D-maltoside. Blots were stained against ATP synthase α subunit to detect F_1_ intermediates

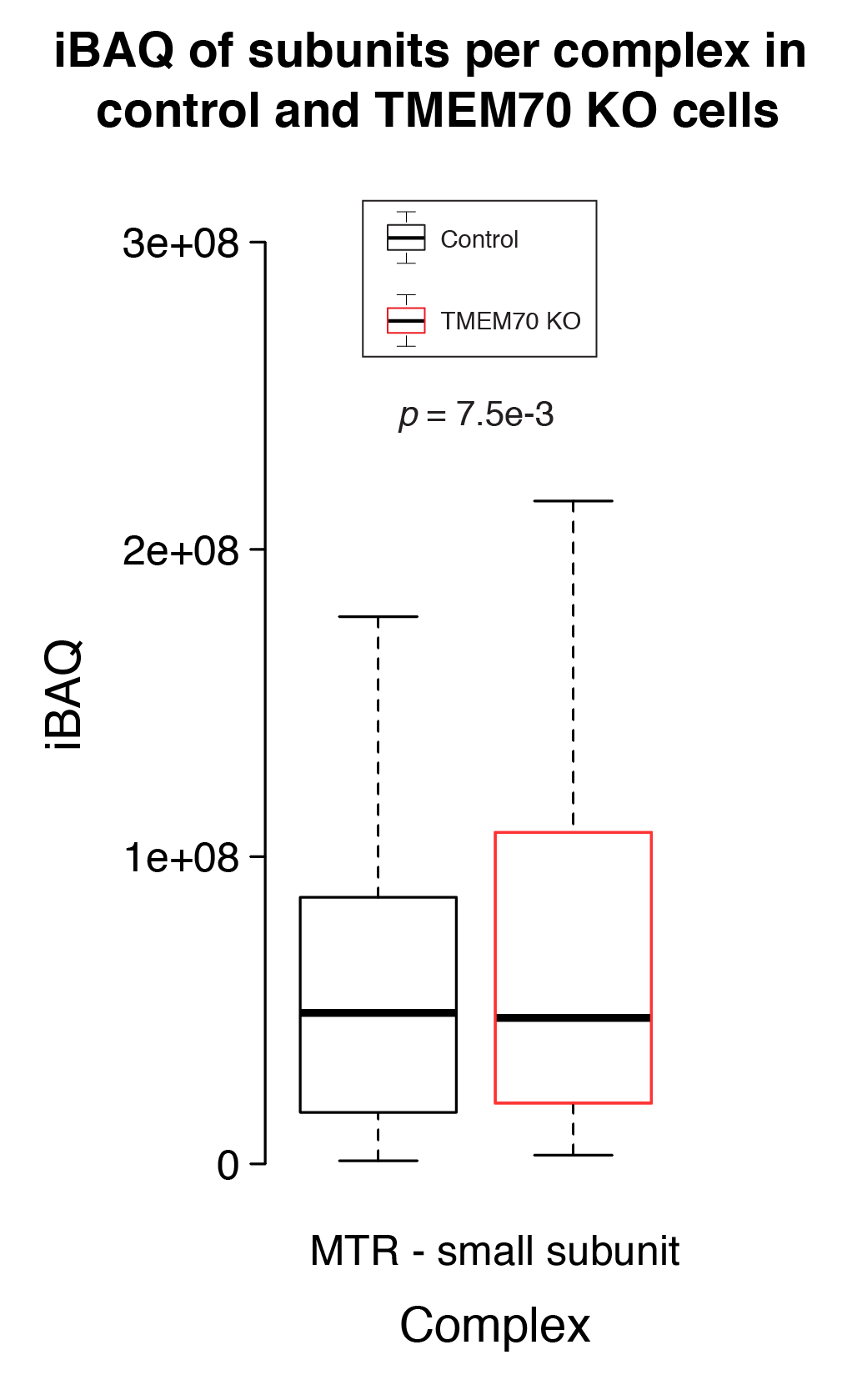

Figure S9. Boxplot depicting the distribution of the iBAQ values of the proteins belonging to the small subunit of the mitochondrial ribosome at the mass of the fully assembled complex.

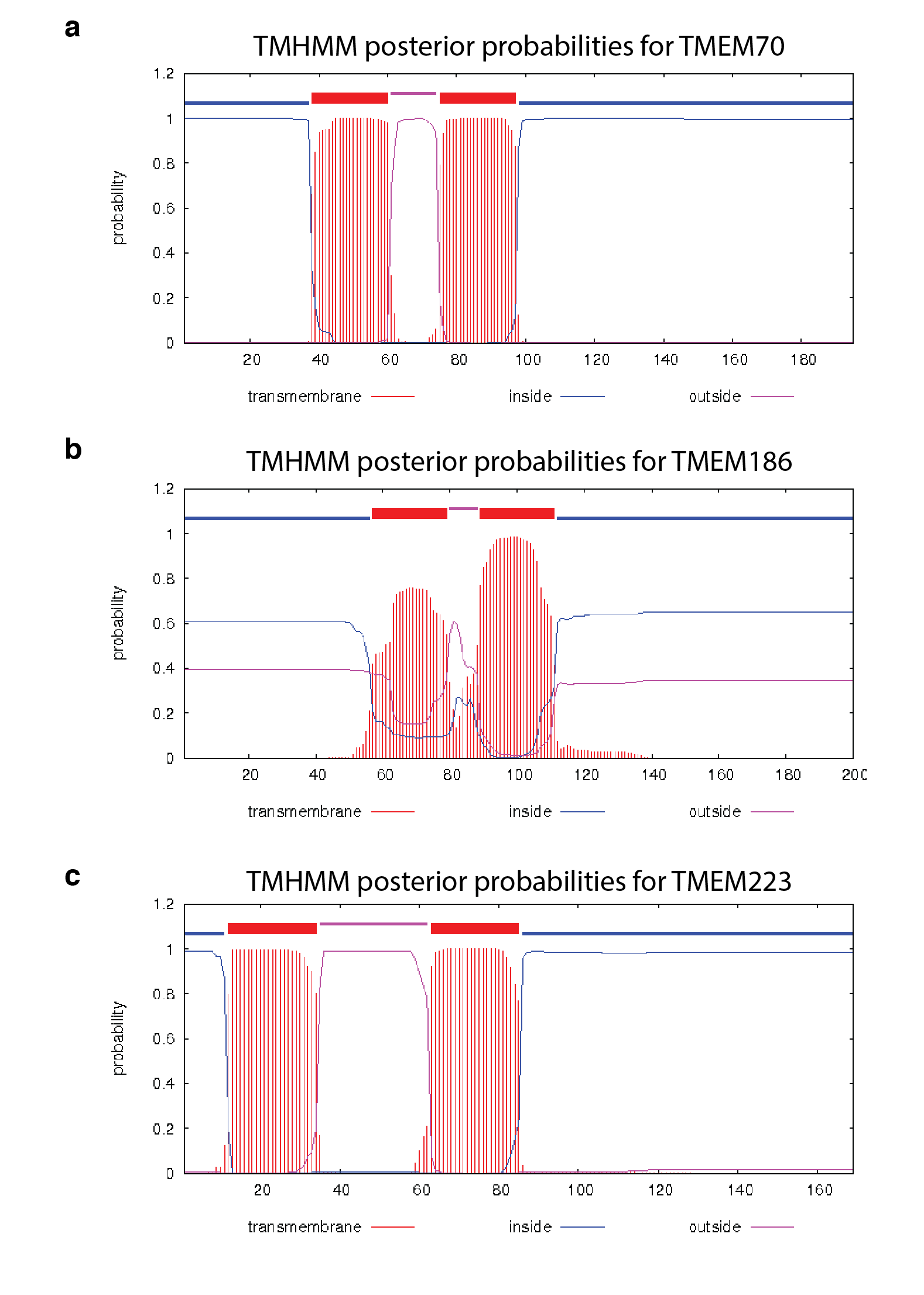

Figure S10. Topology predictions calculated with TMHMM^58^ for **a** TMEM70, **b** TMEM186 and **c** TMEM223 show a similar in-out-in topology with two transmembrane domains close to the N-terminal.

Table S1. iBAQ values of the different BioID experiments. dP: doxycycline positive. dN: doxycycline negative. bP: biotin positive. bN: biotin negative

Table S2. Additional information about the proteins considered as potential interactors of TMEM70

Table S3. Output of DAVID after analysing the potential interactors of TMEM70 using Gene Ontology categories.

Table S4. Normalized iBAQ values obtained by complexome profiling.

Table S5. Normalized iBAQ values obtained by complexome profiling after chymotrypsin digestion

Table S6. Top co-evolving mitochondrial proteins with TMEM70. Mitochondrial proteins sorted by similarity of their phylogenetic profiles compared to TMEM70 as determined by differential Dollo parsimony. The orthologous group ids indicate to which OrthoMCL based orthology a protein belongs to, and thus on which orthologous group the phylogenetic profile is based. Proteins with the same orthologous group id are considered in-paralogs and thus share the same phylogenetic profile.

Table S7. Phylogenetic distribution of TMEM70, TMEM186 and TMEM223. Separation of the family into the various orthologous groups has been done using the phylogeny (Fig. 5), the ciliate (*T. thermophila and P. tetraurelia*) members of the TMEM70/TMEM186/TMEM223 protein family could not confidently be assigned to any specific group however, and where therefore left blank. Phylogenetic distribution of complex I, V and TMEM14 has been established using reciprocal best hits established with Blast searches and, when homologs were not detectable using the Blast, using profile based searches^59^.
